# Control of Golgi- V-ATPase through Sac1-dependent co-regulation of PI(4)P and cholesterol

**DOI:** 10.1101/2024.11.04.621832

**Authors:** Xin Zhou, Miesje M. van der Stoel, Shreyas Kaptan, Haoran Li, Shiqian Li, Maarit Hölttä, Helena Vihinen, Eija Jokitalo, Christoph Thiele, Olli Pietiläinen, Shin Morioka, Junko Sasaki, Takehiko Sasaki, Ilpo Vattulainen, Elina Ikonen

## Abstract

Sac1 is a conserved phosphoinositide phosphatase, whose loss-of-function compromises cell and organism viability. Here, we employed acute auxin-inducible Sac1 degradation to identify its immediate downstream effectors in human cells. Most of Sac1 was degraded in ∼1 h, paralleled by increased PI(4)P and decreased cholesterol in the trans-Golgi network (TGN) during the following hour, and superseded by Golgi fragmentation, impaired glycosylation, and selective degradation of TGN proteins by ∼4 h. The TGN disintegration resulted from its acute deacidification caused by disassembly of the Golgi V-ATPase. Mechanistically, Sac1 mediated TGN membrane composition maintained an assembly promoting conformation of the V_0_a2 subunit. Key phenotypes of acute Sac1 degradation were recapitulated in human differentiated trophoblasts, causing processing defects of chorionic gonadotropin, in line with loss-of-function intolerance of the human *SACML1* gene. Collectively, our findings reveal that the assembly of the Golgi V-ATPase is controlled by the TGN membrane via Sac1 fuelled lipid exchange.

## Introduction

PI(4)P is the most abundant species of phosphoinositides that serve as major regulators of organelle identity and dynamics^1^. It is asymmetrically distributed in the cell, with high levels in the trans-Golgi network (TGN) and the plasma membrane (PM)^2^, regulated by an interplay between organelle specific PI4-kinases (PI4Ks) and the endoplasmic reticulum (ER)/Golgi integral phosphatase Sac1^3–5^. In the Golgi, PI(4)P plays an important role by controlling the localization, signalling and activity of PI(4)P binding proteins and subsequently lipid homeostasis and vesicular trafficking^6–11^. Importantly, PI(4)P in the TGN serves as counter-exchange cargo for lipid transfer reactions at membrane contact sites. Here, PI(4)P regulates the docking of lipid transfer proteins, such as oxysterol binding protein (OSBP), which transfers PI(4)P from the TGN to the ER and in exchange relocates cholesterol to the opposite direction, from the ER to the TGN^12–14^. At the ER, Sac1 hydrolyzes PI(4)P into phosphatidylinositol (PI), thereby creating an intracellular PI(4)P gradient that drives anterograde cholesterol transport and reduces PI(4)P levels in the TGN^13,15,16^. Besides OSBP, other lipid transporters such as ORP9 and Asters/GRAMDs, are also involved in maintaining the cholesterol distribution between the ER and TGN^17^. Sac1 depletion causes an increase in PI(4)P^10,18^ and results in enlargement of the Golgi and mislocalization of glycosylation enzymes^19,20^. Additionally, by locally controlling Golgi PI(4)P levels, Sac1 can regulate cargo trafficking and secretion^20,21^. However, how Sac1 mediated cholesterol and PI(4)P distribution affects Golgi functioning and which critical effectors are involved in this, are currently unknown.

Sac1, or suppressor of actin-1, was first discovered in yeast, where it regulates actin cytoskeletal organization and Golgi secretion^22,23^. Sac1 is the only known PI(4)P phosphatase in mammalian cells and its catalytic domain is highly conserved across species^24^. Constitutive loss of Sac1 compromises the viability of cells and organisms, causing early embryonic lethality in e.g. *Drosophila* and mice^19,25^. As long-term loss of Sac1 compromises viability, the precise mechanisms of how Sac1 fuelled lipid exchange affects the function of the Golgi have been challenging to study. Indeed, due to the rapid turnover and broad distribution of phosphoinositide species in cellular membranes, most if not all membrane compartments may become afflicted in chronic Sac1 depletion models.

Here, we employed auxin-inducible degradation (AID) of Sac1 to rapidly deplete the protein in several human cell types, allowing us to investigate early effects after Sac1 loss. We identified vacuolar-type ATPase (V-ATPase) as a key effector of Sac1 controlled PI(4)P and cholesterol content in the TGN membrane, impacting TGN pH and resulting in defects in Golgi terminal glycosylation and secretion, with physiological implications for early embryonic development. Mechanistically, increased PI(4)P levels in the TGN characteristic of Sac1 loss, promoted a conformational change of the V_0_a2 subunit of V-ATPase preventing the assembly of the Golgi V-ATPase holo-complex. Taken together, this study reveals the Golgi V-ATPase as a critical effector of Sac1 fuelled PI(4)P/cholesterol exchange to regulate Golgi integrity and cargo processing.

## Results

### Sac1 exerts acute control of Golgi morphology

To investigate the acute consequences of Sac1 depletion on the Golgi, we took advantage of the AID system that we recently established in several human cell types^26,27^ (**Fig. 1a**). Within 1 h after IAA addition, about 90% of endogenous Sac1 was removed from A431 cells (**Fig. 1b**). At this time, cellular PI(4)P was already moderately increased, as assessed by mass spectrometry^28^, and after 2 h of IAA treatment, the PI(4)P content was robustly elevated, to ∼3-fold from the starting level (**Fig. 1c**). Instead, no significant or very minor changes were observed in PI(3)P, PI(4,5)P_2_, and PI(3,5)P_2_ species at this time (**Suppl. Fig. 1a-c**), highlighting PI(4)P as the primary substrate of Sac1 in cells. All acyl chain compositions of PI(4)P species were similarly affected, whereas no changes in the acyl chain composition of the other PIP species were observed (**Suppl. Fig. 1d**). By 4 h of Sac1 depletion, all detected phosphoinositide species were elevated compared to the starting situation (**Suppl. Fig. 1e-h**).

**Figure 1:**
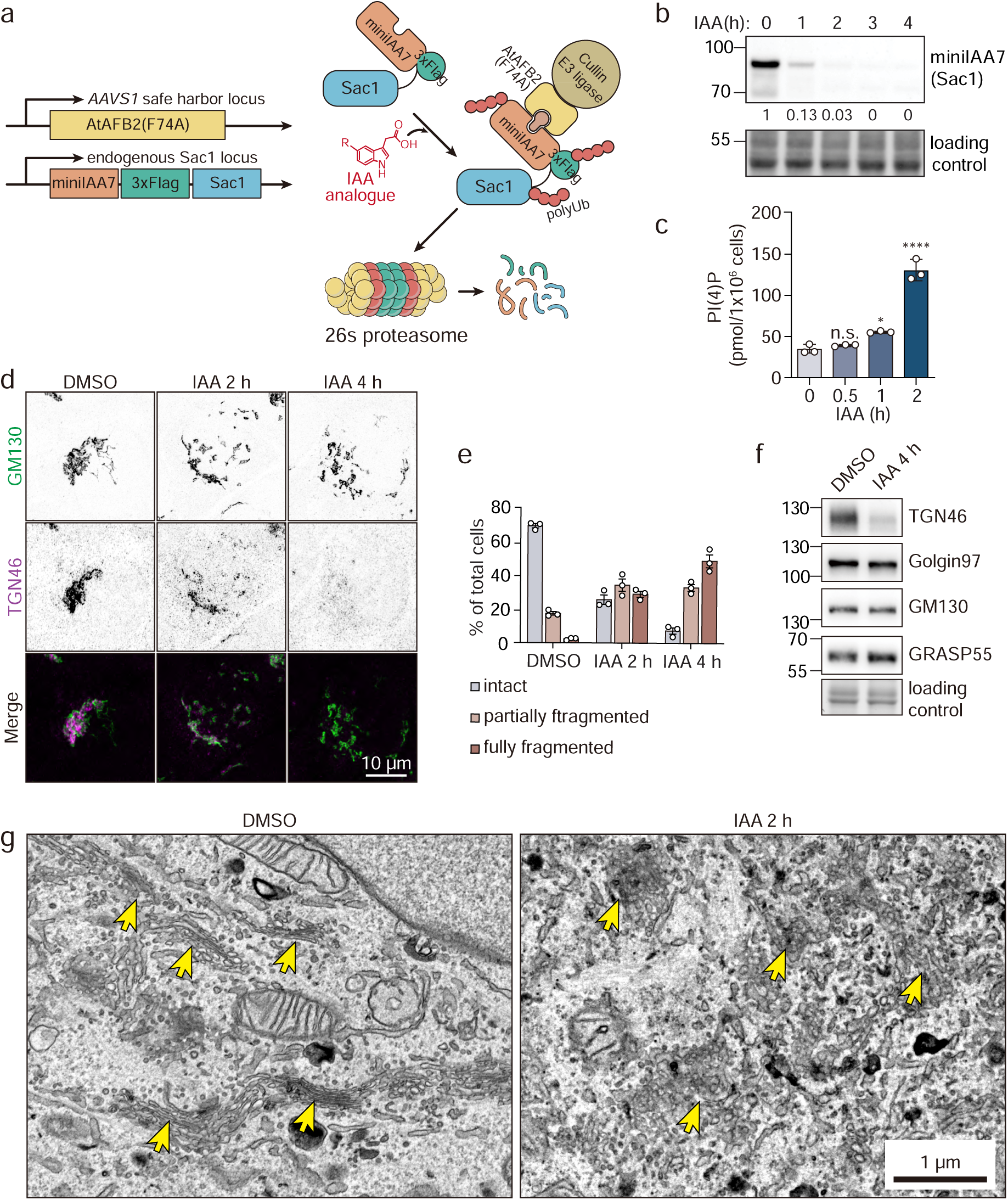
Rapid Sac1 depletion disturbs Golgi morphology. **a)** Schematic overview of the genetic manipulations for auxin inducible degradation of Sac1. Sac1 is endogenously tagged with a miniIAA7-3xFLAG tag and the AtAFB2(F74A)-Cullin E3 ligase complex is exogenously expressed in cells. After addition of an IAA analogue, Sac1-miniIAA7- 3xFLAG is recruited to the AtAFB2(F74A)-Cullin E3 ligase complex, resulting in polyubiquitination and proteasomal degradation of Sac1. **b)** Western blot analysis of Sac1-degron cell lysates after IAA addition for 0 to 4 h in A431 cells, blotted for miniIAA7 to detect Sac1. Normalized relative intensity of each band is marked under the blot. **c)** Bar graph showing total PI(4)P levels ±S.E.M. (pmol/x10^6^ cells) of Sac1-degron A431 cells treated with DMSO for 2 h (0) or IAA for 0.5, 1 or 2 h as measured by PRMC-MS. n = 3 biological replicates. One-way ANOVA with Tukey’s multiple comparisons test. n.s. = non-significant, * p < 0.05, **** p < 0.001. **d)** Representative confocal images of Sac1-degron A431 cells treated with DMSO (4 h) or IAA for 2 or 4 h and stained for GM130 (cis-Golgi, green) or TGN46 (trans-Golgi, magenta). **e)** Bar graph depicting the percentage (mean ±S.E.M) of cells with intact, partially fragmented, or fully fragmented Golgi. Data from 3 independent experiments. **f)** Western blot analysis of Sac1-degron A431 cells treated with DMSO or IAA for 4 h and blotted for TGN46, Golgin-97, GM130 and GRASP55. **g)** Representative TEM images of Sac1-degron A431 cells treated with DMSO or IAA for 2 h. The yellow arrows indicate the Golgi stacks.

Previously, Sac1 depletion was shown to affect Golgi morphology^10,19–21^. To investigate the effect of Sac1 degradation on Golgi integrity, we performed immunofluorescence staining using Golgi markers. After 2 h of IAA induction, the normally tight perinuclear clustering of cis- and trans-Golgi elements (GM130 and TGN46 as markers, respectively) was lost and the ratio of cells with aberrantly fragmented Golgi increased (**Fig. 1d, e**). By 4 h, GM130-positive structures were largely fragmented and TGN46 was hardly detectable by fluorescence microscopy (**Fig. 1d, e).** This was reflected by reduced TGN46 protein levels, while the protein levels of Golgin-97 (trans), GRASP55 (medial) and GM130 (cis) remained unchanged (**Fig. 1f**).

Consistent with these results, thin-section transmission electron microscopy (TEM) revealed a rapid transformation of Golgi ribbons to tubular-vesicular networks in 2 h of IAA induction (**Fig. 1g**). While the Golgi had a typical ribbon-like organization in 80% of the DMSO treated cells, the Golgi elements were dispersed in 70% of the Sac1 degraded cells: the occurrence and length of stacks decreased, and the Golgi elements were mainly compromised of tubular networks and tubulo-vesicular clusters (**Fig. 1g, Extended Data 1**). Importantly, the rapid fragmentation of the Golgi upon Sac1 removal was also evident in other Sac1-degron models, such as HEK293 and A549 cells (**Suppl. Fig. 1i, j**). These results show that Sac1 is important to maintain Golgi integrity.

### Sac1 phosphatase activity is required for terminal glycosylation

To obtain an unbiased overview of the effects of acute Sac1 removal at the protein level, we performed quantitative proteomic analysis of A431 Sac1-degron cells. After 3 h of IAA induction, a total of 8040 proteins were identified, of which 203 were upregulated (fold change ≥ 1.3, IAA to DMSO) and 103 downregulated (fold change ≤ 0.77, IAA to DMSO) (**Suppl. Fig. 2a** and **Extended Data 2**). Among the most upregulated proteins in the Sac1-degron cell proteomics were PI(4)P binding proteins, such as OSBP, ORP9 and CERT (**Suppl. Fig. 2a**). Gene ontology (GO) analysis revealed that the most downregulated proteins were enriched in the biological processes and cellular components related to glycosylation, Golgi organization and membrane (**Fig. 2a** and **Extended Data 2**). Interestingly, while the levels of ER and cis-medial-Golgi glycosylation enzymes were unaltered or moderately downregulated, the enzymes of the trans-Golgi cisternae, in particular galactosyl- and sialyltransferase, were strongly downregulated (**Fig. 2b**).

**Figure 2:**
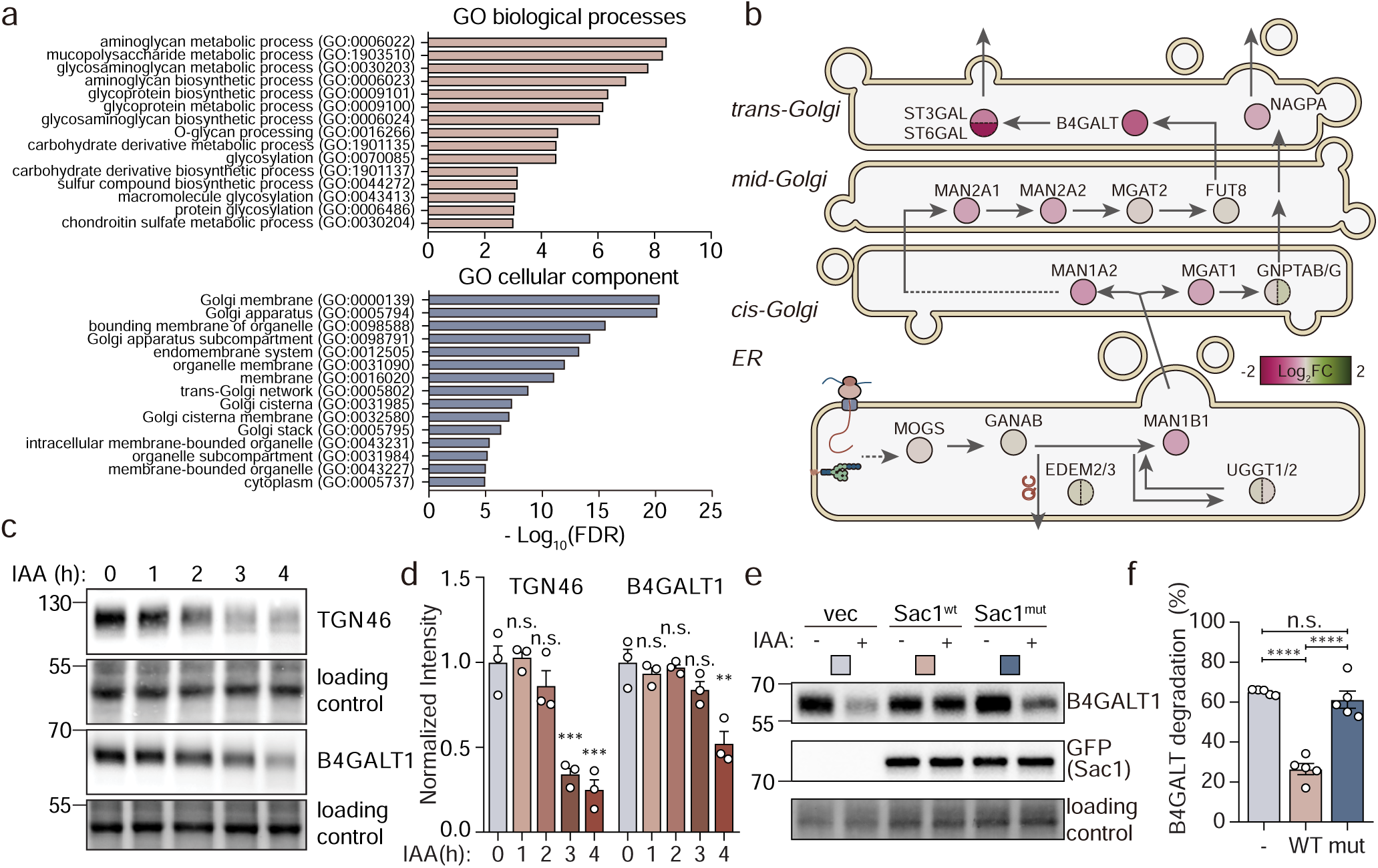
Terminal glycosylation is impaired by Sac1 depletion. **a)** Top terms in gene ontology enrichment analysis of the downregulated proteins in Sac1-degron A431 cells treated with IAA for 3 h compared to DMSO control (-Log_10_ False Discovery Rate (FDR) with a fold change threshold of ≥ 1.30 or ≤ 0.77 and an adjusted p-value of ≤ 0.05). **b)** Schematic overview illustrating the changes in protein levels of glycosylation enzymes in Sac1- degron cells after 3 h IAA treatment. The Log_2_ fold change (Log_2_FC) of individual enzyme is presented using a color gradient from magenta (-2) to green (+2). Glycosylation starts in the ER, where precursor glycan chains are added to the protein. The glycans are modified sequentially by glycosidases and glycosyltransferases along the ER and Golgi stacks. **c)** Western blot analysis of B4GALT1 and TGN46 in Sac1-degron A431 cells treated with IAA for 0 to 4 h. **d)** Bar graph showing normalized intensity of B4GALT1 and TGN46 quantified from **(c)**. n = 3 independent experiments. One-way ANOVA with Dunnett’s multiple comparisons test. n.s. = non- significant, ** p < 0.01, *** p<0.005. **e)** Western blot analysis of lysates of HEK293A Sac1-degron cells transfected with pcDNA3.1 (empty vector), EGFP-Sac1-WT (Sac1^wt^) or EGFP-Sac1-C389S (Sac1^mut^), treated for 6 h with DMSO or IAA and blotted for B4GALT1 and GFP (Sac1). **j)** Graph depicts the B4GALT1 degradation efficiency (mean ±S.E.M) in Sac1-degron HEK293A cells treated for 6 h with IAA and rescued with Sac1^wt^ or Sac1^mut^. n = 5 independent experiments. One-way ANOVA with Tukey’s multiple comparisons test. n.s. = non-significant, **** p<0.001.

Western blotting confirmed the robust reduction of beta-1,4-galactosyltransferase 1 (B4GALT1) and of TGN46 (**Fig. 2c, d**) by 4 h of IAA induction, while the mRNA levels of B4GALT1 and TGN46 remained unchanged (**Suppl. Fig 2b**). To investigate if lysosomal degradation plays a role in the loss of TGN46, we neutralized lysosomes using Bafilomycin A1 (BafA1) during IAA treatment. This effectively restored TGN46 protein levels (**Suppl. Fig. 2c**), suggesting that TGN46 undergoes lysosomal proteolysis upon Sac1 depletion. Interestingly, at 8 h the mature form of TGN46 was replaced by an immature form of the protein (**Suppl. Fig. 2d**), and PNGaseF and α-2-3,6,8,9- Neuraminidase A treatments indicated defects in its terminal glycosylation (**Suppl. Fig. 2e, f**).

To scrutinize if the effect of Sac1 on TGN integrity was dependent on its phosphatase activity, we reintroduced EGFP-tagged wild-type Sac1 (Sac1^wt^) and catalytically inactive mutant Sac1-C389S (Sac1^mut^) to Sac1 degraded cells, using the well transfectable HEK293A Sac1-degron cells. After 24 h of transient transfection, endogenously degron-tagged Sac1 was removed by 6 h of IAA induction, while the levels of exogenous Sac1^wt^ and Sac1^mut^ remained stable (**Fig. 2e**). In this setting, the reduction of B4GALT1 levels caused by Sac1 degradation could be rescued by reintroduction of Sac1^wt^ but not by Sac1^mut^ (**Fig. 2e, f**). Collectively, these results suggest that Sac1 mediated PI(4)P hydrolysis maintains TGN integrity and terminal glycosylation.

### Sac1 is needed to maintain TGN pH and Golgi V-ATPase assembly status

The Golgi maintains a proton gradient, from neutral pH at the cis face to pH ∼6 at the TGN. This is important to maintain Golgi integrity and protein glycosylation^29^. To assess if acute Sac1 degradation affects the pH of the TGN, we transfected Sac1-degron cells with TGN targeting GalT-RpHLuorin2, a pH-sensitive GFP mutant fused to B4GALT1^30^ (**Suppl. Fig. 3a**). Fluorescence lifetime values of GalT-RpHLuorin2 were obtained from A431 Sac1-degron cells and converted to pH values according to a standard curve (**Suppl. Fig. 3b**).

The average pH of the TGN of A431 cells remained consistently ∼5.9 in the DMSO control group over time, while it was elevated to ∼6.1 at 2 h and ∼6.3 at 3 h of IAA induction (**Fig. 3a** and **Suppl. Fig. 3c**). However, when we applied EN6, an activator of vacuole-type ATPase (V-ATPase)^31^ to counteract the pH elevation, the TGN pH dropped to ∼6.0 in Sac1-depleted cells (**Fig. 3b**). Although EN6 marginally affected inducible Sac1 depletion, re-acidification of the TGN efficiently prevented Sac1 depletion induced B4GALT1 and TGN46 degradation (**Fig. 3c, d**), and partially restored the TGN morphology (**Fig. 3e, f**). These data argue that acute Sac1 depletion deacidifies the TGN, thereby inducing selective TGN morphological changes and degradation.

**Figure 3:**
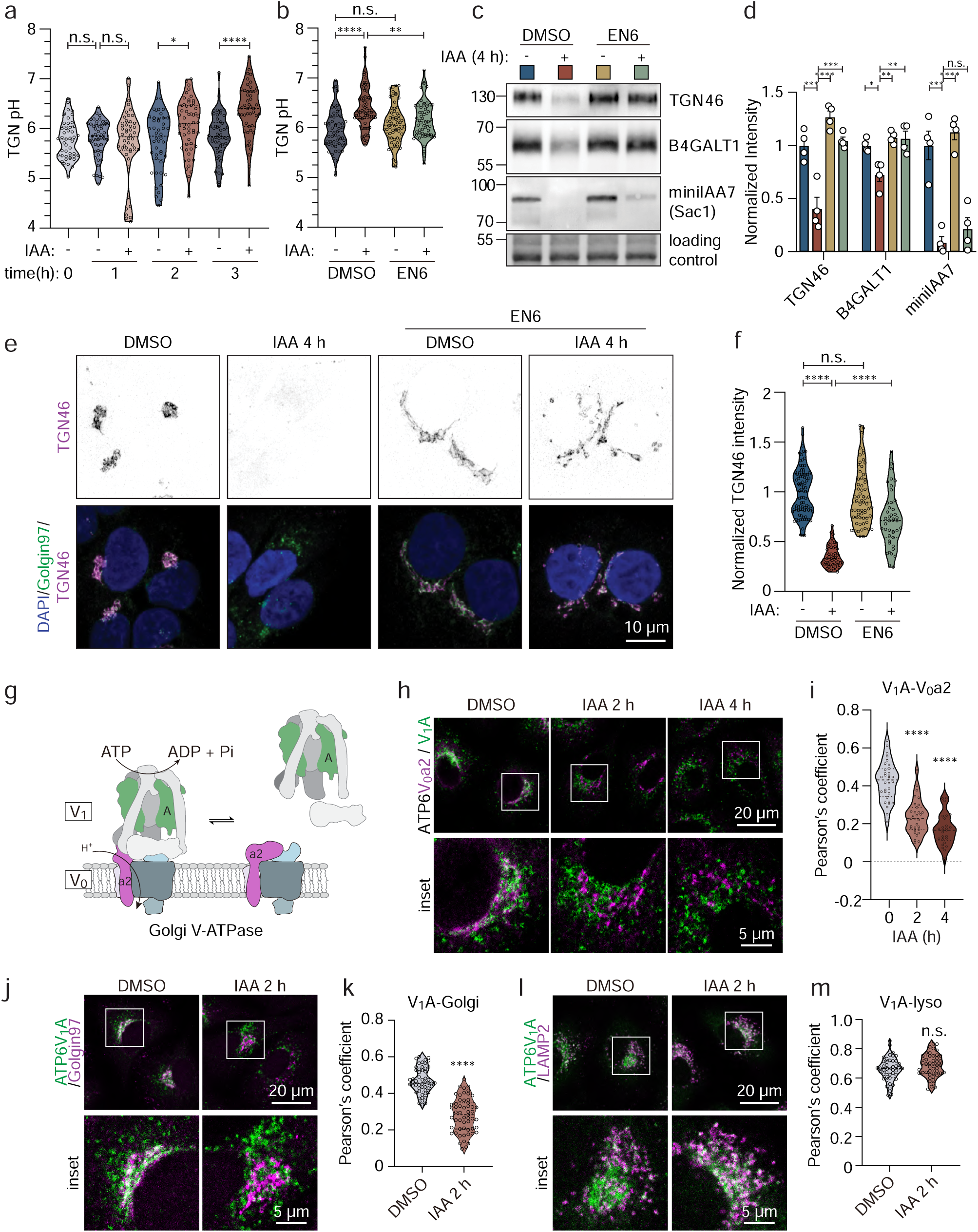
Acute Sac1 depletion results in TGN deacidification caused by V-ATPase disassembly. a) Violin plot showing the distribution of TGN pH values of Sac1-degron A431 cells transiently transfected with GalT-RpHLuorin2 and treated with DMSO or IAA for 0 to 3 h. Data from 3 independent experiments, n = 46 – 54 cells. One-way ANOVA with Tukey’s multiple comparisons test. n.s. = non-significant, * p<0.05, **** p<0.001. b) Violin plot depicting the distribution of TGN pH values of Sac1-degron A431 cells transiently transfected with GalT-RpHLuorin2 treated for 3 h with DMSO or IAA with or without 50 µM EN6. Data from 3 independent experiments, n = 55 – 63 cells. One-way ANOVA with Tukey’s multiple comparison test. n.s. = non-significant, ** p<0.01, **** p<0.001. c) Western blot analysis of Sac1-degron A431 cells treated for 4 h with DMSO or IAA with or without 50 µM EN6. Immunoblotted for TGN46, B4GALT1 and miniIAA7 (Sac1) with a representative stain free loading control. d) Bar graph showing average normalized immunoblot intensities from **(c)**. Data from 4 independent experiments. One-way ANOVA with Tukey’s multiple comparisons test. n.s. = non-significant, * p<0.05, ** p<0.01, *** p<0.005, **** p<0.001. e) Representative confocal images of Sac1-degron A431 cells treated for 4 h with DMSO or IAA with or without 50 µM EN6. Stained for DAPI (blue), Golgin-97 (green) and TGN46 (magenta). f) Violin plot depicts the distribution of the TGN46 intensity in the Golgi area calculated from **(e)**. Data from 3 independent experiments, n = 42 – 84 cells. One-way ANOVA with Dunnet’s multiple comparisons test. n.s. = non-significant, **** p<0.001. g) Cartoon illustrating the reversible assembly of the V_0_ and V_1_ regions of the V-ATPase complex. Upon assembly, ATP hydrolysis of the V_1_ region drives the proton pumping ability of the V_0_ region. h) Representative fluorescence micrographs of Sac1-degron A431 cells treated with DMSO for 4 h or IAA for 2 or 4 h and stained for ATP6V_0_a2 (magenta) or ATP6V_1_A (green). i) Violin plot showing the distribution of the Pearson’s correlation coefficient between ATP6V_0_a2 and ATP6V_1_A from (**h**). Representative data from one of three different experiments. One-way ANOVA with Tukey’s multiple comparisons test. **** p<0.001. j) Representative confocal images of Sac1-degron A549 cells treated with DMSO or IAA for 2 h and stained for ATP6V_1_A (green) and Golgin-97 (magenta). k) Violin plot showing the Pearson’s correlation coefficient between ATP6V_1_A and Golgin-97 from **(j)**. n = 53 – 55 cells. Student’s t-test. **** p<0.001. l) Representative confocal images of Sac1-degron A549 cells treated with DMSO or IAA for 2 h and stained for ATP6V_1_A (green) and LAMP2 (magenta). m) Violin plot showing the Pearson’s correlation coefficient between ATP6V_1_A and LAMP2 from **l)**. n = 45 – 46 cells. Student’s t-test. n.s. = non-significant.

V-ATPase is a ubiquitous and essential regulator of organelle pH, with the cytoplasmic V_1_ region consuming ATP to facilitate proton pumping through the membrane-embedded V_0_ region. The reversible assembly and disassembly of the V_0_ and V_1_ regions orchestrates proton translocation to adapt to various physiological demands (**Fig. 3g**)^32^. We therefore studied if the V-ATPase complex is affected by Sac1 depletion. The total amounts of V_0_- and V_1_-subunits were not altered by acute Sac1 degradation, as assessed by specific antibodies (**Suppl. Fig. 3d**). However, membrane-cytosol fractionation suggested that upon 2 h of Sac1 degradation the membrane-association of the V_1_A subunit was reduced (**Suppl. Fig 3e, f**).

We next studied the subcellular localization of the V_1_ region. For this, we employed Sac1-degron A549 cells, due to the prominent cytoplasmic V_1_A immunoreactivity in A431 cells that complicated the assessment. We found substantial colocalization of the V_1_A subunit with the Golgi-specific V_0_a2 subunit in control conditions, as expected^33^. Remarkably, this colocalization diminished gradually during 2-4 h after inducing Sac1 degradation (**Fig. 3h, i**). Co-staining with Golgin-97 (**Fig. 3j, k**) and LAMP2 (**Fig. 3l, m**) antibodies revealed that at 2 h after inducing Sac1 degradation, the TGN localization of the V_1_ region was reduced, while its lysosomal localization was maintained. In parallel, the Golgi specific V_0_a2 subunit clearly overlapped with the Golgi marker GM130, while no colocalization with the lysosomal marker LAMP2 was observed (**Suppl. Fig. 3g**). Together, these data suggest that the V_0_ and V_1_ regions of the Golgi V-ATPase complex become disassembled rapidly after Sac1 depletion, providing a mechanism for the deacidification of the TGN.

### Sac1 controls V-ATPase complex integrity via its lipid environment

As Sac1 fuels PI(4)P/cholesterol lipid exchange by LTPs, i.e. OSPB^13,14,17^, we assessed the role of Sac1 in controlling PI(4)P and cholesterol distribution in the TGN. Consistently with the mass spectrometry results (**Fig. 1c**), expression of mCherry-P4M-SidM and immunostaining with PI(4)P antibody in A431 Sac1-degron cells revealed an increasing PI(4)P signal over time, starting perinuclearly at early time points after degradation and extending later on throughout the cell (**Suppl. Fig. 4a-c**). The latter agrees with findings from chronic Sac1 depletion systems^10,18,21^. Remarkably, already at 1 h of Sac1 degradation, we observed a prominent accumulation of PI(4)P in the TGN region (**Fig. 4a, b**). Staining of live cells with mNeonGreen-ALOD4 recognizing PM accessible cholesterol^34^, showed that after 1 h of Sac1 removal, the PM accessible cholesterol levels were slightly reduced, and by 4 h, only ∼20% of control levels remained **(Suppl. Fig. 4d, e**). Filipin staining and thin-layer chromatography showed a minor reduction in total cellular free cholesterol over time (**Suppl. Fig. 4f-h**). However, by 2 h after Sac1 degradation, the filipin signal in the TGN area was significantly reduced (**Fig 4c, d**) and in the lysosomes slightly increased (**Suppl. Fig. 4i, j)**. Meanwhile, cholesterol esterification was increased 2.5-fold by 2 h (**Suppl**. **Fig. 4k**) and total cholesteryl ester content elevated 1.5-fold by 4 h (**Suppl. Fig. 4l**), in line with increased cholesterol delivery to the ER. Together, these results provide evidence that degradation of Sac1 rapidly alters both the PI(4)P and cholesterol content of the TGN.

**Figure 4:**
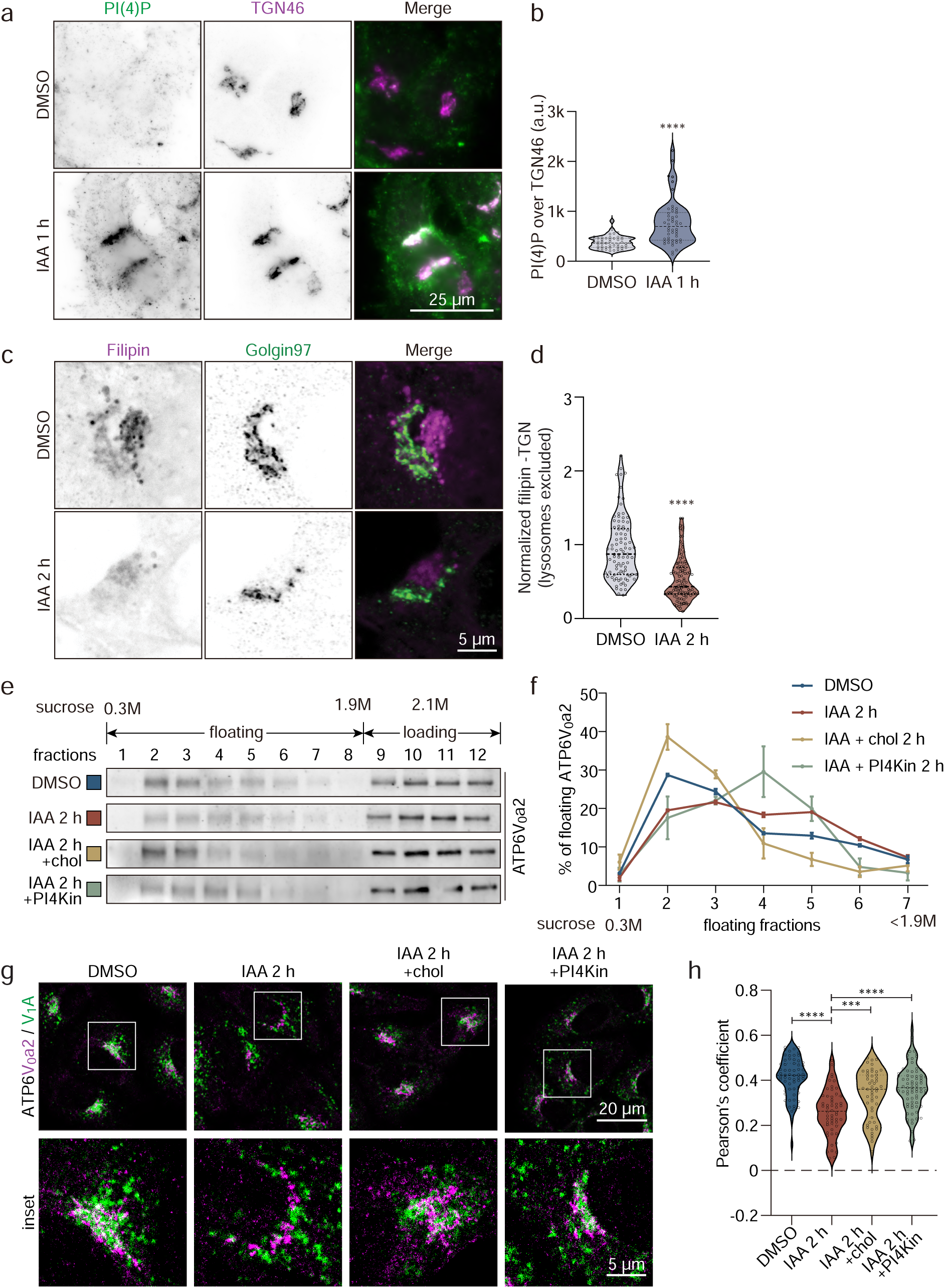
Sac1 fuelled PI(4)P/cholesterol exchange maintains V-ATPase activity. a) Representative confocal images of Sac1-degron A431 cells treated with IAA or DMSO for 1 h and stained for PI(4)P (green) and TGN46 (magenta). b) Violin plot showing the average PI(4)P intensity in the Golgi area (TGN46). Representative data from one of three independent experiments. n = 50 cells. Student’s t-test. **** p<0.001. c) Representative confocal images of Sac1-degron A431 cell treated with IAA or DMSO for 2 h and stained with Filipin (magenta) and Golgin-97 (green). d) Violin plot showing the normalized filipin intensity in the Golgi area (Golgin-97). Data from 3 independent experiments. n = 92 – 107 cells. Student’s t-test. **** p<0.001. e) Western blot analysis of sucrose gradient fractionations (Sucrose concentration from 0.3M (fraction 1) to 2.1M (loading fractions 9 – 12)) of A549 Sac1-degron cells treated for 2 h with DMSO, IAA, IAA with cholesterol, or IAA with PI4K inhibitors (NC03 and PIK93) and blotted for ATP6V_0_a2. f) Line graph depicting the average percentage ±S.E.M. of floating ATP6V_0_a2 distributed between fraction 1 to 7 of A549 Sac1-degron cells treated for 2 h with DMSO, IAA, IAA plus cholesterol, or IAA plus PI4K inhibitors (NC03 and PIK93). n = 3 technical replicates. g) Representative confocal images of A549 Sac1-degron cells treated for 2 h with DMSO, IAA, IAA in combination with cholesterol or PI4K inhibitors. Stained for ATP6V_1_A (green) and ATP6V_0_a2 (magenta). h) Violin plot depicting the distribution of the Pearson’s correlation coefficient between ATP6V_1_A and ATP6V_0_a2 from (**g**). Data from one of three independent experiments. One-way ANOVA with Tukey’s multiple comparisons tests. *** p<0.005, **** p<0.001.

Next, we investigated if the changes in the TGN lipid environment induced by Sac1 degradation affect the Golgi V-ATPase by using sucrose density gradient fractionation of cellular post-nuclear supernatants (PNS). We found that the floating Golgi V-ATPase V_0_ region peaked in the light membrane fraction 2, as probed by anti-V_0_a2 antibody (**Fig. 4e, f**). Upon Sac1 depletion for 2 h, the flotation of the V_0_ region became less efficient, suggesting that the membrane environment of the V_0_ region shifted towards higher equilibrium density (**Fig. 4e, f**). Interestingly, cellular cholesterol depletion phenocopied the effect of Sac1 depletion on V_0_ flotation (**Suppl. Fig. 5a**), suggesting that the cholesterol reduction in the TGN membrane contributes to the behavior of the V_0_ region in Sac1 depleted cells.

To probe this more directly, we replenished cells with cholesterol while Sac1 was being degraded. This effectively rescued the flotation of the V_0_ region (**Fig. 4e, f**). We also tested the effect of inhibiting PI(4)P generation by PI(4)P kinase (PI4K) inhibitors (PIK93 for PI4KIIIβ and NC03 for PI4K2A), the main PI4Ks in the Golgi^6^, during Sac1 degradation. This did not revert the V_0_ region to low-density membrane fractions (**Fig. 4e, f**), suggesting that PI(4)P, as a minor membrane constituent, does not affect the overall flotation behavior of the V_0_ region. Together, these findings suggest that cholesterol reduction in the TGN induced by Sac1 removal affects the TGN membrane environment of the V_0_ region.

Next, we analyzed if cholesterol supplementation or PI4K inhibitor treatment during Sac1 degradation can rescue V_0_-V_1_ complex assembly. Interestingly, both increasing membrane cholesterol or decreasing PI(4)P generation improved the V_0_-V_1_ colocalization in Sac1 depleted cells (**Fig. 4g, h**), while only PI4K inhibition could partially rescue Golgi morphology (**Suppl**. **Fig. 5b-d**). Collectively, the data suggest that Sac1 degradation alters the TGN membrane lipid composition resulting in V-ATPase complex disassembly, providing a mechanism for the TGN deacidification and glycosylation defects.

### The conformation of the V_0_ region is influenced by the membrane lipid composition

To investigate how the membrane lipid composition affects Golgi V-ATPase, we carried out atomic- level molecular dynamics simulations of the V_0_ region containing the Golgi specific V_0_a2 subunit (**Fig. 5a**), focusing on the role of PI(4)P and cholesterol (**Fig. 5b**, **Table 1**). We analyzed the simulation data using deep learning techniques, looking for structural classes where the V_0_ region conformations are similar. Using this approach, the conformational space was first reduced to two dimensions using an autoencoder, after which these states were clustered with a Bayesian Gaussian Mixture Model, searching for structural classes representing distinct conformations. The analysis revealed four separate clusters (**Fig. 5c**) characterized by balanced populations (**Suppl. Fig. 6a**) and high structural similarity within each cluster (**Suppl. Fig. 6b**). A representative conformation of the V_0_ region from each of the four clusters is depicted in **Fig. 5d** (top view) and **Suppl. Fig. 6c** (side view). Based on structural data of the V_0_-V_1_ complex, the assembly of the V_0_ and V_1_ regions requires the detachment of the terminus (a2 head) of the V_0_a2 subunit from the d1 subunit, because otherwise the a2 head sterically hinders V_0_-V_1_ complex assembly. Similar structural rearrangements of the yeast V_0_a1 N-terminal domain have been described to be essential for V_0_-V_1_ assembly/disassembly^35^.

**Figure 5:**
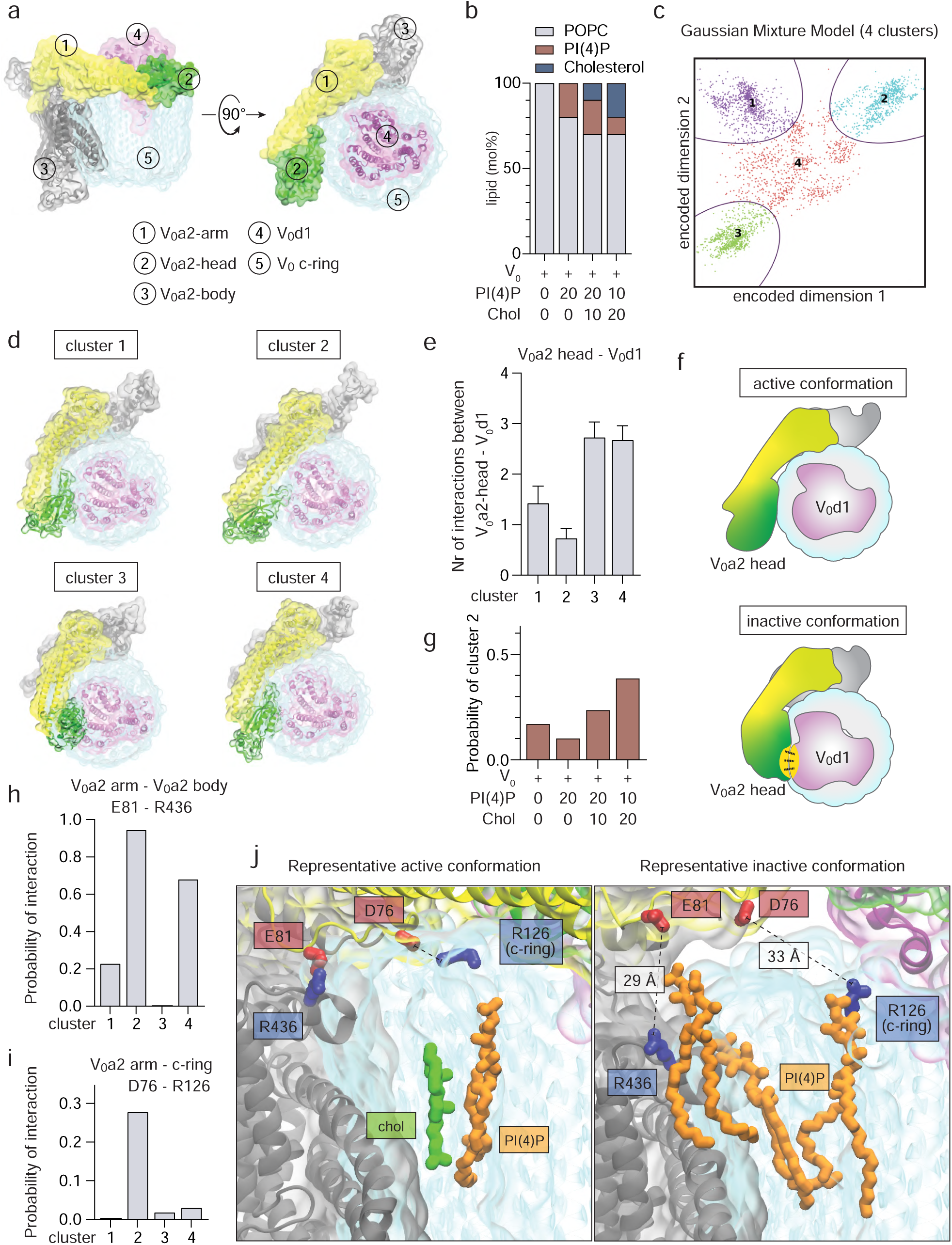
Atomic-level molecular dynamics simulations of the V_0_ region in different lipid membrane compositions. a) Top and side view of the Golgi V-ATPase V_0_ region, highlighting different subunits. The V_0_a2 subunit is divided into three regions: a2 body (grey), a2 arm (yellow), and a2 head (green). The d1 subunit is shown in magenta. The c-ring is shown in cyan. b) Graphical representation of the different membrane lipid compositions used in the simulations. c) Clustering of the simulation data using Bayesian Gaussian Mixture Models (BGMM) reveals four distinct clusters, each representing a different structural class. The clusters are divided by their decision boundaries, which represent the region where the uncertainty of a cluster label approaches 50%. d) Representative top view of the V_0_ region from each of the four clusters. For each cluster, the structure with the smallest distance from the mean of the Gaussian in the BGMM is presented. Cluster 2 has the a2 head (green) furthest from the d1 subunit (magenta). e) Average number of salt bridge interactions formed by the V_0_a2-head with the V_0_d1 subunit. The error bars are the standard error computed over 32 independent datasets composed of 8 independent replicas with 4 different lipid compositions. f) Schematic representation of the active conformation (cluster 2) and the inactive conformations, where the V_0_a2 head interacts with the V_0_d1 subunit g) Probability (relative population) of cluster 2 with each membrane lipid composition. h) Probability of forming the salt bridge E81 (V_0_a2 arm) – R436 (V_0_a2 body) in the four different clusters. i) Probability of forming the salt bridge D76 (V_0_a2 arm) – R126 (V_0_ c-ring) in the four different clusters. j) Representative images of active and inactive conformations of the V_0_ region showing how the lipids (PI(4)P in orange, cholesterol in green) affect the formation of the D76 (V_0_a2-arm) – R126 (c-ring) and E81 (V_0_a2-arm) – R436 (V_0_a2-body) salt bridges. The dotted line between the residues indicates the distance between the residues, a salt bridge is defined when the residues lie within 4.0 Å of each other^97^. Of note, the two salt bridges cannot form simultaneously, therefore the representative active conformation presents a conformation where the involved residues are closest together.

Analysis of the simulation data showed that among the four conformational clusters detected, cluster 2 plays a crucial role in the assembly process, as in these conformations the a2 head formed the fewest interactions with the d1 subunit (**Fig. 5e**). Thus, cluster 2 characterizes conformations of the V_0_ region that are prone to bind to the V_1_ region to form a functional V_0_-V_1_ complex. Notably, these conformations differ from those in the other clusters (**Fig. 5f**). PLS-based discriminant analysis of the simulation data (**Movie 1, 2**) showed that, compared to the other clusters, in cluster 2 the arm and head of the a2 subunit are positioned close to the membrane and do not sterically interfere with the binding orientation of the V_1_ region.

### High level of PI(4)P suppresses the conformational state of the V_0_ region that promotes V_0_ -V_1_ complex assembly

We next analyzed the probability of occurrence of cluster 2 in membranes of different lipid compositions (**Fig. 5g**). We found that the cluster 2 population is minimized (<10% of conformations) when a cholesterol-free membrane is enriched with ample (20 mol%) PI(4)P. When 10 mol% cholesterol is added to this membrane, the population of cluster 2 increases to ∼24%. The population of cluster 2 is at its highest (40%) when the concentration of PI(4)P is reduced to 10 mol% and in parallel cholesterol is increased to 20 mol%. These data imply that in the absence of cholesterol, high PI(4)P suppresses the assembly-promoting conformational state of V_0_ region, while cholesterol plays an assembly-promoting, rescuing role.

### Cholesterol blocks the access of PI(4)P deep into the pocket between the a2-subunit and the c-ring

From the atomic-level simulation data, we elucidated the mechanisms of PI(4)P and cholesterol action by focusing on their densities in the immediate vicinity of the V_0_ region (**Suppl. Fig. 6d**). In membranes with high PI(4)P and without cholesterol (mimicking Sac1 degradation), we found that PI(4)P in the cytosolic membrane leaflet interacts with the a2 arm, a2 body, and the c-ring, disrupting two potential salt bridges in the pocket between the a2 subunit and the c-ring. These salt bridges are between E81 of the a2 arm and R436 of the a2 body (**Fig. 5h**) and between D76 of the a2 arm and R126 of the c-ring (**Fig. 5i, j**), stabilizing the orientation of the a2 head away from the d1 subunit. Of these, the salt bridge E81-R436 is present in several clusters but most prominent in cluster 2, with a probability of ∼90% (**Fig. 5h**), while the salt bridge D76 - R126 practically only forms in cluster 2 and has a probability of 30% (**Fig. 5i**). Instead, in cholesterol and PI(4)P containing membranes promoting V_0_-V_1_ complex assembly, cholesterol replaces PI(4)P binding on the cytosolic side of the membrane in the pocket between the a2 and the c-ring (**Fig. 5j, Suppl. Fig. 6d**). This suggests that cholesterol prevents the disruption of the salt bridges between the a2 arm, the a2 body and the c-ring by competing with PI(4)P for these interactions.

### Sac1 safeguards hCG processing in human trophoblasts

Our experimental findings identified an essential role for Sac1 in maintaining Golgi morphology and terminal glycosylation via V-ATPase mediated pH regulation. We next challenged the physiological relevance of this idea. As Sac1 loss in mouse is embryonically lethal^19,25^, variations in the *SACM1L* gene might not be well tolerated in humans. Indeed, based on the Genome Aggregation Database (gnomAD) the probability of the *SACM1L* gene for being loss-of-function intolerant (lof.pLI) is high, 0.97 (**Fig. 6a**). Additionally, the observed number of loss-of-function *SACM1L* variants is only 27% of the expected (o/e = 0.27, 90% CI: 0.15-0.51), consistent with negative selection acting on these variants in humans and with an essential role of Sac1 also in human development.

**Figure 6:**
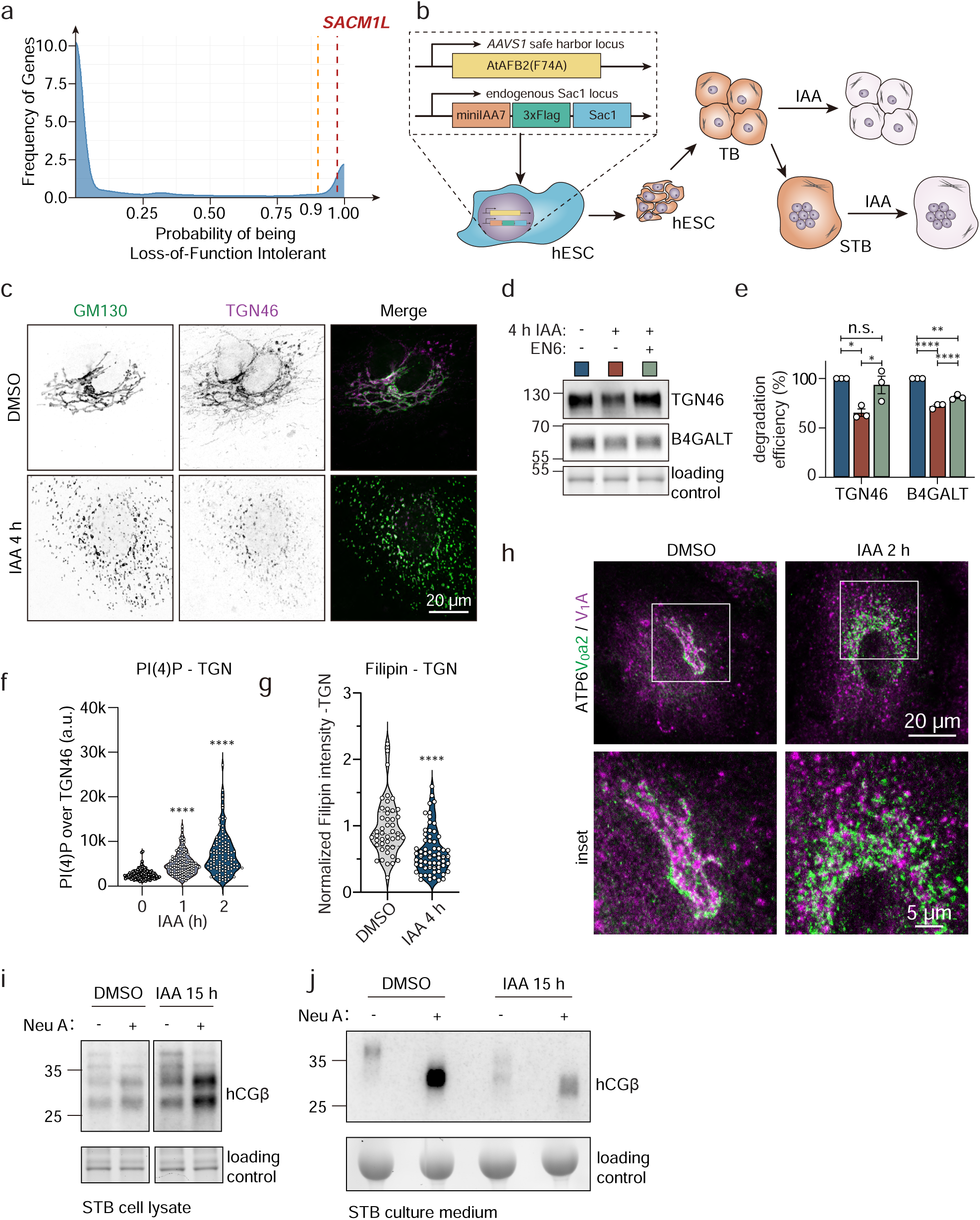
Sac1 depletion impairs hCG secretion in trophoblasts. **a)** Density plot of gnomAD data set showing the frequency of genes (y-axis) of having the probability of being loss of function intolerant (lof.pLI, x-axis). The orange dotted line indicates the 0.9 threshold commonly used for Mendelian diseases and the red dotted line the location of the *SACM1L* gene. **b)** Schematic overview of differentiation of trophoblasts from Sac1-degron hESCs. Sac1-degron hESCs are subsequently differentiated into trophoblasts (TB) and syncytiotrophoblasts (STB). IAA treatment can induce Sac1 degradation in both TB and STB. **c)** Representative confocal images of Sac1-degron STB treated for 4 h with IAA and stained for GM130 (green) and TGN46 (magenta). **d)** Western blot analysis of Sac1-degron STB treated for 4 h with DMSO or IAA and 50 µM EN6 and blotted for TGN46 and B4GALT1. **e)** Bar graph depicting the TGN46 or B4GALT1 degradation efficiency calculated from (**d**). Data from 3 biological replicates. One-way ANOVA with Tukey’s multiple comparisons test. n.s. = non- significant, * p<0.05, **** p<0.001. **f)** Violin plot showing the distribution of average PI(4)P staining intensity in the TGN. n > 100 cells. One-way ANOVA with Dunnett’s multiple comparisons test. **** p<0.001. **g)** Violin plot showing normalized filipin intensity in the TGN area (excluding lysosomal signal). Data from 3 independent experiments. n = 46 – 63 cells. Mann-Whitney U test. **** p<0.001. **h)** Representative confocal images of Sac1-degron STBs treated for 2h with DMSO or IAA, stained for ATP6V_0_a2 (green) and ATP6V_1_A (magenta). **i)** Western blot analysis of cell lysate and secreted medium of Sac1-degron STB treated with DMSO or IAA for 15 h, with or without Neuraminidase A digestion and blotted for hCGβ. **j)** Western blot analysis of secreted medium of Sac1-degron STBs treated with DMSO, IAA or IAA for 15 h, with or without Neuraminidase A digestion. Blotted for hCGβ.

During the early stages of mammalian embryogenesis, secretion of growth factors, hormones and cytokines orchestrates close communication between the blastocyst and the uterine wall. Particularly, human chorionic gonadotropin (hCG), a trophoblast-secreted glycoprotein, is required for implantation and gestation^36^. To investigate if Sac1 is important for hCG secretion by trophoblasts, we differentiated human embryonic stem cells (ESCs) carrying the Sac1-degron system^27^ into trophoblasts (**Fig. 6b**). After 10 days of differentiation, the trophoblast markers hCGβ (beta-subunit of hCG), GATA3 and Dab2 were readily detected (**Suppl Fig. 7a**), and multi-nucleated syncytiotrophoblasts were obtained after an additional 6 days of differentiation. Importantly, the levels of hCGβ increased drastically during differentiation, as anticipated (**Suppl. Fig. 7b**).

Acute depletion of Sac1 from trophoblasts led to a severe fragmentation of the Golgi by 4 h (**Fig. 6c**), followed by reduced protein levels of TGN46 and B4GALT1 (**Suppl. Fig. 7c, d**). Remarkably, when EN6 was added to the trophoblasts together with IAA, the TGN46 protein levels could be restored; B4GALT1 levels were also partially rescued (**Fig. 6d, e**). At 1-2 h of Sac1 degradation, PI(4)P levels rapidly increased in the TGN (**Fig. 6f, Suppl. Fig. 7e**), while TGN cholesterol levels decreased (**Fig. 6g, Suppl. Fig. 7f, g**). Sac1 degradation in trophoblasts also acutely induced V-ATPase disassembly within 2 h (**Fig 6h**).

Next, we assessed the impact of Sac1 depletion on trophoblast hCGβ processing and secretion. At 15 h upon Sac1 removal, the cellular levels of hCGβ were increased and the highly glycosylated hCGβ, abundantly secreted in normal conditions, was massively reduced in the medium (**Fig. 6i, j**). Neuraminidase treatment of hCGβ increased the antibody reactivity and showed that hCGβ was abundantly sialylated in control trophoblasts and that some residual sialylation took place in Sac1 depleted cells. However, upon Sac1 degradation, hCGβ with a lower molecular weight was secreted, implicating a glycosylation defect (**Fig. 6j**). Overall, this demonstrates that key phenotypes observed upon acute Sac1 degradation are recapitulated in differentiated trophoblasts and highlights the Golgi V-ATPase complex as a key effector of Sac1 activity that safeguards hCGβ processing and secretion in trophoblasts.

## Discussion

This study employs the potential of AID in elucidating the early functional consequences of eliminating a vital protein, Sac1. The rapid, tightly controlled degradation of Sac1 enabled us to systematically follow the time course of ensuing defects, thereby revealing underlying causalities. In approximately an hour, Sac1 was fully degraded, followed by a ∼3-fold increase in PI(4)P concentrated in the TGN area and a decrease in free cholesterol levels in the TGN during the following hour. These changes in the TGN lipid environment prevent the assembly of the Golgi V-ATPase complex as observed from 2 h onwards, subsequently elevating TGN pH from ∼5.9 to ∼6.3 and resulting in disruption of terminal glycosylation and selective TGN degradation by 4 h (**Fig. 7**). Importantly, these phenotypes could be recapitulated in human differentiated trophoblasts, resulting in defects in hCGβ glycosylation and secretion. This may in part explain the preimplantation lethality of Sac1 null mice and rationalizes the intolerance for Sac1 loss-of-function during mammalian development.

**Fig 7.**
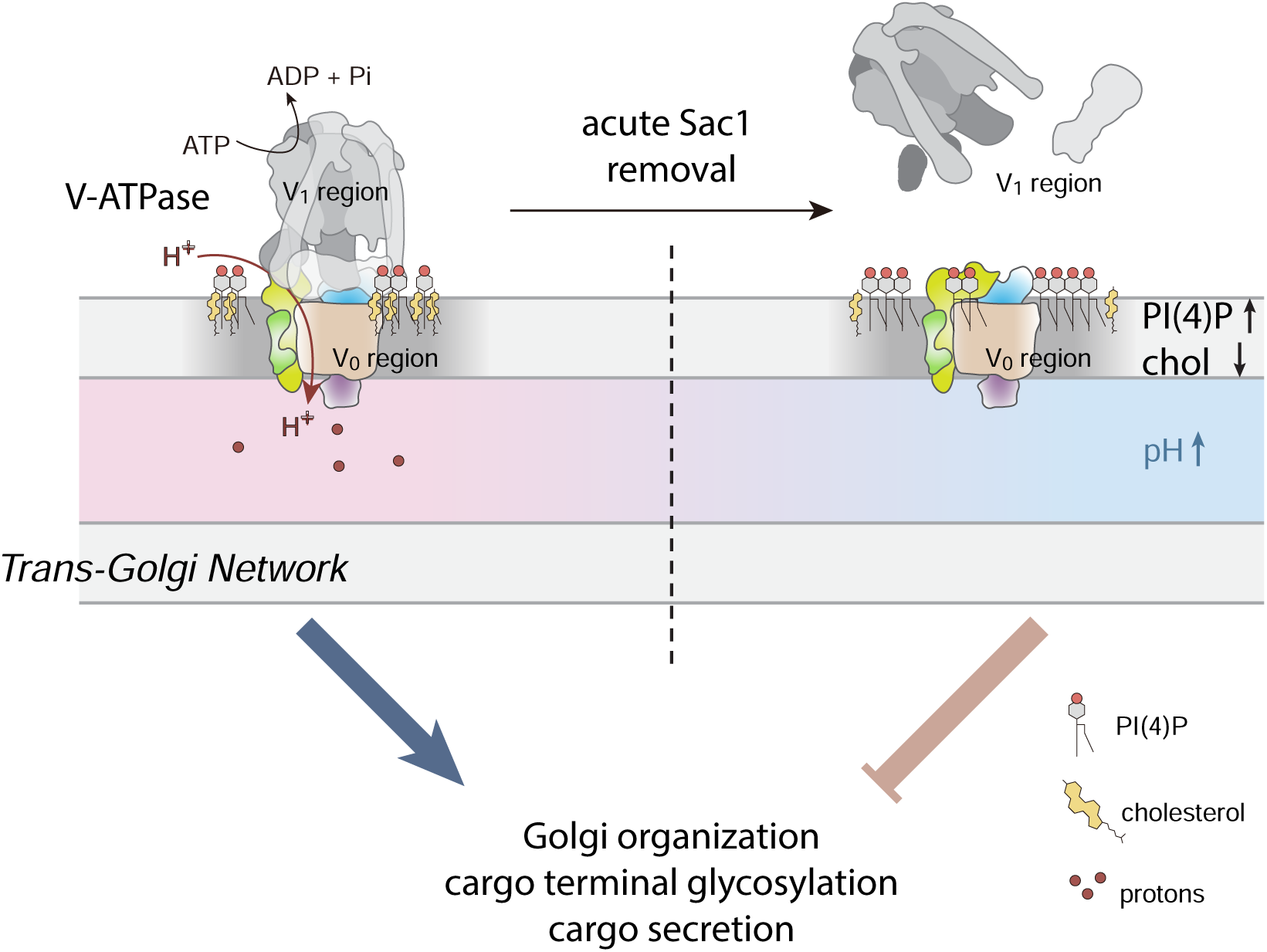
Working model for Sac1 controlled TGN V-ATPase activity and downstream effects. Sac1 fuelled PI(4)P/cholesterol exchange maintains the composition of the Golgi membrane, allowing V-ATPase assembly for TGN acidification, proper terminal glycosylation and cargo secretion. Rapidly after Sac1 degradation, PI(4)P is accumulated in the TGN while cholesterol is depleted. This changes the lipid microenvironment of the V_0_ region, resulting in a conformational change in the V_0_a2 subunit which prevents the assembly of the Golgi V-ATPase complex. This results in the deacidification of the TGN, Golgi fragmentation and defects in cargo processing and secretion.

The most direct effect of loss of Sac1 function is the accumulation of its ligand PI(4)P that facilitates cargo trafficking from the TGN ^10,11,18,37^. Importantly, we provide evidence that Sac1 is needed not only to maintain PI(4)P levels, but also the high cholesterol content in the TGN, in line with the critical role of Sac1 in fuelling PI(4)P/cholesterol exchange by OSBP at ER-TGN contact sites^11,13,14,38^. As complex and intricate interactions between lipid transfer proteins control cholesterol and PI(4)P levels in the TGN^17^, Sac1 most likely affects a variety of lipid transfer proteins beyond OSBP. The upregulation of several lipid transporters, observed in our proteomics data, probably serves to compensate for the acute loss of Sac1. However, this is clearly insufficient to maintain the normal ER-TGN (and ER-plasma membrane) cholesterol gradient without Sac1 burning off PI(4)P. Hence, the critical role of Sac1 in governing cellular cholesterol distribution contributes to its vital importance.

The present study highlights the TGN membrane lipid dependent control of Golgi V-ATPase activity via Sac1 as a shared denominator and driver for the later phenotypes, i.e. Golgi deacidification, fragmentation and degradation, as well as cargo glycosylation and secretion defects. This notion is supported by the observations that TGN fragmentation, protein degradation and defective cargo secretion could be partly rescued by pharmacological re-acidification. It also agrees with the established role of V-ATPase in safeguarding Golgi glycosylation and secretion^39–44^. As Sac1 activity has also been implicated in cargo secretion from the TGN^20,21,45^, the coupling of TGN carrier formation to V-ATPase activity may help to coordinate terminal glycosylation and secretion.

V-ATPases are key regulators of organelle acidification conserved from plants to humans. The regulation of V-ATPases has been best characterized in *Saccharomyces cerevisiae*, where the association/dissociation of V_0_ and V_1_ regions rapidly adjusts proton pumping activity, with e.g. glucose starvation stimulating complex disassembly and conservation of energy^46^. In mammalian cells, the cues controlling the reversible association of V_0_ and V_1_ regions may be more context dependent, with e.g. glucose removal promoting either assembly or dissociation^32,47^, and studies have mainly focused on the lysosomal V-ATPase^48–51^.

Despite V-ATPases being large integral membrane protein complexes, there is limited understanding on how they are functionally regulated by lipids. In yeast, V-ATPase activity can be modulated by phosphoinositides, phospholipids and ergosterol^33,52–55^. The human a-subunits of V_0_ were shown to bind to distinct phosphoinositides, with the endo-lysosomal isoform binding to PI(3)P and PI(3,5)P_2_ and the Golgi isoform to PI(4)P *in vitro*^33,56,57^. Importantly, the cryo-EM structures of human V- ATPase reveal several lipids, including phospholipids and cholesterol, as integral parts of the V_0_ region^58,59^.

The present study complements and extends these findings by identifying Sac1 as a gatekeeper whose activity controls, via effects on the TGN membrane lipid environment, the conformation and activity of Golgi V-ATPase in the cellular context. Specifically, we provide evidence that the dissociation of the V_0_ and V_1_ regions is promoted by increased PI(4)P and decreased cholesterol in the TGN membrane upon Sac1 removal. Atomic-level simulations illuminate the mechanistic role of cholesterol in stabilizing the assembly-promoting conformation of the V_0_ region. In the absence of cholesterol, PI(4)P can interact with residues forming salt bridges in the recess between the a2 subunit and the c-ring of the V_0_ region and thus destabilize the assembly-promoting conformation. The assembly-disrupting conformations lead to binding of the a2 head to the d1 subunit, which sterically prevents the V_1_ region from assembling with the V_0_ region. Cholesterol, on the other hand, prevents PI(4)P from entering deep into this recess, thereby promoting assembly.

Based on our data, the V-ATPase assembly status was more sensitive to loss of Sac1 function at Golgi than lysosomal membranes: upon 2 h of Sac1 removal - when PI(4)P was the only phosphoinositide increased – the V_1_ region became disassembled from the Golgi V_0_ region but the localization of the V_1_ region in lysosomal membranes appeared intact. Moreover, at 4 h of Sac1 removal, TGN46 degradation was still sensitive to lysosomal V_0_-V_1_ dissociation by BafA1. The early affliction of Golgi V-ATPase may be related to different phosphoinositide sensitivities of the V-ATPases, but possibly also to a predominant activity of Sac1 at Golgi contacts.

The initial effects of Sac1 removal that we focused on, centred on the TGN. Yet, other compartments, such as endo-lysosomes, are likely affected over time. Interestingly, recent data implicate lysosomal PI(4)P, generated by local synthesis and turned over by Sac1, in lysosomal repair^60,61^ and rewiring of lysosomal degradation during starvation^62^. We found all the major PIPs to be elevated by 4 h of Sac1 removal. As PIPs are rapidly hydrolyzed and phosphorylated by specific kinases and phosphatases, the initial increase of PI(4)P is expected to result in the interconversion to other PIP species over time^63^. Thus, a broad PIP imbalance affecting multiple endomembranes and PIP dependent lipid transfer processes may contribute to the long-term effects of Sac1 depletion. Moreover, considering the coupling between PIP and cholesterol distribution, it is likely that cholesterol and PIPs co-operate in the regulation of membrane proteins, including V-ATPase, in other compartments.

In conclusion, this study reveals that in human cells, the Golgi V-ATPase assembly status is dynamically modulated by membrane lipids, specifically by the TGN PI(4)P and cholesterol pools that are physiologically co-regulated by Sac1 activity. The multiple modes of lipid-V-ATPase interactions and how they regulate V-ATPase activity will be important topics for future work.

## Acknowledgements

We thank the HiLIFE and Biocenter Finland supported Biomedicum Imaging and Electron Microscopy Units and Turku Proteomics Facility for infrastructure support, Giray Enkavi and Waldemar Kulig for discussions on molecular dynamics simulations, and Anna Uro for technical assistance. XZ was supported by Academy of Finland grant 332096, MMvdS and HZ received funding from the European Union’s Horizon 2020 Research and Innovation programmes under the Marie Sklodowska-Curie Personal Fellowship (MMvdS) no. 101059424 and the Marie Sklodowska- Curie Grant Agreement (HZ) no. 953489. EI was supported by the Academy of Finland (grant no. 324929), Foundation Leducq 19CVD04, Jane and Aatos Erkko Foundation and Sigrid Juselius Foundation, TS was supported by AMED (grant no. 24gm1710007), by TMDU under Multilayered Stress Diseases (JPMXP1323015483) and Medical Research Center Initiative for High Depth Omics.

IV thanks the Helsinki Institute of Life Science (HiLIFE) Fellow program, Sigrid Jusélius Foundation, Lundbeck Foundation, and Academy of Finland (project IDs: 335527, 331349, 336234, 364185) for financial support. The authors thank CSC–IT Center for Science Ltd (Espoo, Finland) for providing computational resources.

## Author contributions

XZ and MMvdS contributed equally. EI conceived and oversaw the project. XZ, MMvdS, HL, MH, performed experiments and analyzed data. SL provided Sac1-degron cell lines and technical support for cell engineering. HV and EJ performed electron microscopic analysis. CT performed click-lipid MS experiments. OP performed gnomAD dataset analysis. SM, JS and TS performed PRMC-MS experiments and analysis. SK and IV performed and analyzed atomistic molecular dynamics simulations. XZ, MMvdS and EI wrote the manuscript, and all authors edited the manuscript.

## Competing Interests

The authors declare no competing interests.

## Notes

### Competing Interest Statement

The authors have declared no competing interest.

## References

1. Balla, T. Phosphoinositides: Tiny lipids with giant impact on cell regulation. Physiol. Rev. 93, 1019–1137 (2013).

2. Hammond, G. R. V, Machner, M. P. & Balla, T. A novel probe for phosphatidylinositol 4- phosphate reveals multiple pools beyond the Golgi. J. Cell Biol. 205, 113–126 (2014).

3. Balla, A., Tuymetova, G., Barshishat, M., Geiszt, M. & Balla, T. Characterization of Type II Phosphatidylinositol 4-Kinase Isoforms Reveals Association of the Enzymes with Endosomal Vesicular Compartments*. J. Biol. Chem. 277, 20041–20050 (2002).

4. Tan, J. & Brill, J. A. Cinderella story: PI4P goes from precursor to key signaling molecule. Crit. Rev. Biochem. Mol. Biol. 49, 33–58 (2014).

5. Balla, A. et al. Maintenance of Hormone-sensitive Phosphoinositide Pools in the Plasma Membrane Requires Phosphatidylinositol 4-Kinase IIIα. Mol. Biol. Cell 19, 711–721 (2008).

6. Graham, T. R. & Burd, C. G. Coordination of Golgi functions by phosphatidylinositol 4- kinases. Trends Cell Biol. 21, 113–121 (2011).

7. Lolicato, F., Nickel, W., Haucke, V. & Ebner, M. Phosphoinositide switches in cell physiology - from molecular mechanisms to disease. J. Biol. Chem. (2024) doi:10.1016/j.jbc.2024.105757.

8. Wang, Y. J. et al. Phosphatidylinositol 4 Phosphate Regulates Targeting of Clathrin Adaptor AP-1 Complexes to the Golgi. Cell 114, 299–310 (2003).

9. Dippold, H. C. et al. GOLPH3 Bridges Phosphatidylinositol-4- Phosphate and Actomyosin to Stretch and Shape the Golgi to Promote Budding. Cell 139, 337–351 (2009).

10. Wakana, Y. et al. The ER cholesterol sensor SCAP promotes CARTS biogenesis at ER–Golgi membrane contact sites. J. Cell Biol. 220, (2020).

11. Kovács, D. et al. Lipid exchange at ER–trans-Golgi contact sites governs polarized cargo sorting. J. Cell Biol. 223, e202307051 (2023).

12. Moser von Filseck, J., et al. Phosphatidylserine transport by ORP/Osh proteins is driven by phosphatidylinositol 4-phosphate. Science *(80-.).* **349**, 432 LP – 436 (2015).

13. Mesmin, B. et al. A Four-Step Cycle Driven by PI(4)P Hydrolysis Directs Sterol/PI(4)P Exchange by the ER-Golgi Tether OSBP. Cell 155, 830–843 (2013).

14. Mesmin, B. et al. Sterol transfer, PI4P consumption, and control of membrane lipid order by endogenous OSBP. EMBO J. 36, 3156–3174 (2017).

15. Zewe, J. P., Wills, R. C., Sangappa, S., Goulden, B. D. & Hammond, G. R. V. SAC1 degrades its lipid substrate PtdIns4P in the endoplasmic reticulum to maintain a steep chemical gradient with donor membranes. Elife 7, e35588 (2018).

16. Venditti, R. et al. The activity of Sac1 across ER-TGN contact sites requires the four- phosphate-adaptor-protein-1. J. Cell Biol. 218, 783–797 (2019).

17. Naito, T. et al. Regulation of cellular cholesterol distribution via non-vesicular lipid transport at ER-Golgi contact sites. Nat. Commun. 14, 5867 (2023).

18. Dong, R. et al. Endosome-ER Contacts Control Actin Nucleation and Retromer Function through VAP-Dependent Regulation of PI4P. Cell 166, 408–423 (2016).

19. Liu, Y. et al. The Sac1 Phosphoinositide Phosphatase Regulates Golgi Membrane Morphology and Mitotic Spindle Organization in Mammals. Mol. Biol. Cell 19, 3080–3096 (2008).

20. Cheong, F. Y. et al. Spatial Regulation of Golgi Phosphatidylinositol-4-Phosphate is Required for Enzyme Localization and Glycosylation Fidelity. Traffic 11, 1180–1190 (2010).

21. Blagoveshchenskaya, A. et al. Integration of Golgi trafficking and growth factor signaling by the lipid phosphatase SAC1. J. Cell Biol. 180, 803–812 (2008).

22. Novick, P., Osmond, B. C. & Botstein, D. Suppressors of yeast actin mutations. Genetics 121, 659–674 (1989).

23. Cleves, A. E., Novick, P. J. & Bankaitis, V. A. Mutations in the SAC1 gene suppress defects in yeast Golgi and yeast actin function. J. Cell Biol. 109, 2939–2950 (1989).

24. Liu, Y., Boukhelifa, M., Tribble, E. & Bankaitis, V. A. Functional studies of the mammalian Sac1 phosphoinositide phosphatase. Adv. Enzyme Regul. 49, 75–86 (2009).

25. Wei, H.-C. et al. The Sac1 Lipid Phosphatase Regulates Cell Shape Change and the JNK Cascade during Dorsal Closure in Drosophila. Curr. Biol. 13, 1882–1887 (2003).

26. Li, S., Prasanna, X., Salo, V. T., Vattulainen, I. & Ikonen, E. An efficient auxin-inducible degron system with low basal degradation in human cells. Nat. Methods 16, 866–869 (2019).

27. Li, S. et al. HiHo-AID2: boosting homozygous knock-in efficiency enables robust generation of human auxin-inducible degron cells. Genome Biol. 25, 58 (2024).

28. Morioka, S. et al. A mass spectrometric method for in-depth profiling of phosphoinositide regioisomers and their disease-associated regulation. Nat. Commun. 13, 1–9 (2022).

29. Rivinoja, A., Hassinen, A., Kokkonen, N., Kauppila, A. & Kellokumpu, S. Elevated Golgi pH impairs terminal N-glycosylation by inducing mislocalization of Golgi glycosyltransferases. J. Cell. Physiol. 220, 144–154 (2009).

30. Linders, P. T. A., Ioannidis, M., ter Beest, M. & van den Bogaart, G. Fluorescence Lifetime Imaging of pH along the Secretory Pathway. ACS Chem. Biol. 17, 240–251 (2022).

31. Chung, C. Y.-S. et al. Covalent targeting of the vacuolar H(+)-ATPase activates autophagy via mTORC1 inhibition. Nat. Chem. Biol. 15, 776–785 (2019).

32. McGuire, C. M. & Forgac, M. Glucose starvation increases V-ATPase assembly and activity in mammalian cells through AMP kinase and phosphatidylinositide 3-kinase/Akt signaling. J. Biol. Chem. 293, 9113–9123 (2018).

33. Banerjee, S. & Kane, P. M. Direct interaction of the Golgi V-ATPase a-subunit isoform with PI(4)P drives localization of Golgi V-ATPases in yeast. Mol. Biol. Cell 28, 2518–2530 (2017).

34. Infante, R. E. & Radhakrishnan, A. Continuous transport of a small fraction of plasma membrane cholesterol to endoplasmic reticulum regulates total cellular cholesterol. Elife 6, e25466 (2017).

35. Couoh-Cardel, S., Milgrom, E. & Wilkens, S. Affinity Purification and Structural Features of the Yeast Vacuolar ATPase V_o_ Membrane Sector *. J. Biol. Chem. 290, 27959–27971 (2015).

36. Nwabuobi, C. et al. hCG: Biological Functions and Clinical Applications. Int. J. Mol. Sci. 18, (2017).

37. von Blume, J. & Hausser, A. Lipid-dependent coupling of secretory cargo sorting and trafficking at the trans-Golgi network. FEBS Lett. 593, 2412–2427 (2019).

38. Péresse, T., et al. Molecular and cellular dissection of the oxysterol-binding protein cycle through a fluorescent inhibitor. J. Biol. Chem. 295, 4277–4288 (2020).

39. Kornak, U. et al. Impaired glycosylation and cutis laxa caused by mutations in the vesicular H+-ATPase subunit ATP6V0A2. Nat. Genet. 40, 32–34 (2008).

40. Oh-hashi, K., Kanamori, Y., Hirata, Y. & Kiuchi, K. Characterization of V-ATPase inhibitor- induced secretion of cysteine-rich with EGF-like domains 2. Cell Biol. Toxicol. 30, 127–136 (2014).

41. Guillard, M. et al. Vacuolar H+-ATPase meets glycosylation in patients with cutis laxa. Biochim. Biophys. Acta - Mol. Basis Dis. 1792, 903–914 (2009).

42. Serio, M. C. Mutations in V-ATPase assembly factors cause Congenital Disorder of Glycosylation ( CDG) with autophagic liver disease. (2019).

43. Hucthagowder, V. et al. Loss-of-function mutations in ATP6V0A2 impair vesicular trafficking, tropoelastin secretion and cell survival. Hum. Mol. Genet. 18, 2149–2165 (2009).

44. Levic, D. S. et al. Distinct roles for luminal acidification in apical protein sorting and trafficking in zebrafish. J. Cell Biol. 219, e201908225 (2020).

45. Wakana, Y. et al. CARTS biogenesis requires VAP-lipid transfer protein complexes functioning at the endoplasmic reticulum-Golgi interface. Mol. Biol. Cell 26, 4686–4699 (2015).

46. Kane, P. M. Disassembly and reassembly of the yeast vacuolar H(+)-ATPase in vivo. J. Biol. Chem. 270, 17025–17032 (1995).

47. Sautin, Y. Y., Lu, M., Gaugler, A., Zhang, L. & Gluck, S. L. Phosphatidylinositol 3-kinase- mediated effects of glucose on vacuolar H+-ATPase assembly, translocation, and acidification of intracellular compartments in renal epithelial cells. Mol. Cell. Biol. 25, 575–589 (2005).

48. Trombetta, E. S., Ebersold, M., Garrett, W., Pypaert, M. & Mellman, I. Activation of lysosomal function during dendritic cell maturation. Science 299, 1400–1403 (2003).

49. Zoncu, R. et al. mTORC1 Senses Lysosomal Amino Acids Through an Inside-Out Mechanism That Requires the Vacuolar H+-ATPase. Science (80-.). 334, 678–683 (2011).

50. Ratto, E. et al. Direct control of lysosomal catabolic activity by mTORC1 through regulation of V-ATPase assembly. Nat. Commun. 13, 4848 (2022).

51. Hooper, K. M. et al. V-ATPase is a universal regulator of LC3-associated phagocytosis and non-canonical autophagy. J. Cell Biol. 221, e202105112 (2022).

52. Li, S. C. et al. The signaling lipid PI(3,5)P₂ stabilizes V₁-V(o) sector interactions and activates the V-ATPase. Mol. Biol. Cell 25, 1251–1262 (2014).

53. Banerjee, S., Clapp, K., Tarsio, M. & Kane, P. M. Interaction of the late endo-lysosomal lipid PI(3,5)P2 with the Vph1 isoform of yeast V-ATPase increases its activity and cellular stress tolerance. J. Biol. Chem. 294, 9161–9171 (2019).

54. Vasanthakumar, T. et al. Structural comparison of the vacuolar and Golgi V-ATPases from Saccharomyces cerevisiae. Proc. Natl. Acad. Sci. 116, 7272–7277 (2019).

55. Zhang, Y.-Q. et al. Requirement for ergosterol in V-ATPase function underlies antifungal activity of azole drugs. PLoS Pathog. 6, e1000939 (2010).

56. Mitra, C., Winkley, S. & Kane, P. M. Human V-ATPase a-subunit isoforms bind specifically to distinct phosphoinositide phospholipids. J. Biol. Chem. 299, 105473 (2023).

57. Chu, A. et al. Characterization of a PIP Binding Site in the N-Terminal Domain of V-ATPase a4 and Its Role in Plasma Membrane Association. International Journal of Molecular Sciences vol. 24 (2023).

58. Wang, L., Wu, D., Robinson, C. V, Wu, H. & Fu, T.-M. Structures of a Complete Human V- ATPase Reveal Mechanisms of Its Assembly. Mol. Cell 80, 501–511.e3 (2020).

59. Coupland, C. E. et al. High-resolution electron cryomicroscopy of V-ATPase in native synaptic vesicles. Science (80-.). 385, 168–174 (2024).

60. Tan, J. X. & Finkel, T. A phosphoinositide signalling pathway mediates rapid lysosomal repair. Nature (2022) doi:10.1038/s41586-022-05164-4.

61. Radulovic, M. et al. Cholesterol transfer via endoplasmic reticulum contacts mediates lysosome damage repair. EMBO J. 41, e112677 (2022).

62. Ebner, M. et al. Nutrient-regulated control of lysosome function by signaling lipid conversion. Cell 186, 5328–5346.e26 (2023).

63. Hasegawa, J., Strunk, B. S. & Weisman, L. S. PI5P and PI(3,5)P2: Minor, but essential phosphoinositides. Cell Struct. Funct. 42, 49–60 (2017).

64. Wei, Y. et al. Protocol to derive human trophoblast stem cells directly from primed pluripotent stem cells. STAR Protoc. 3, 101638 (2022).

65. Bligh, E. G. & Dyer, W. J. A rapid method of total lipid extraction and purification. Can. J. Biochem. Physiol. 37, 911–917 (1959).

66. Wunderling, K., Zurkovic, J., Zink, F., Kuerschner, L. & Thiele, C. Triglyceride cycling enables modification of stored fatty acids. Nat. Metab. 5, 699–709 (2023).

67. Thiele, C., Wunderling, K. & Leyendecker, P. Multiplexed and single cell tracing of lipid metabolism. Nat. Methods 16, 1123–1130 (2019).

68. Wilhelm, L. P. et al. STARD3 mediates endoplasmic reticulum-to-endosome cholesterol transport at membrane contact sites. EMBO J. 36, 1412–1433 (2017).

69. Jokitalo, E., Cabrera-Poch, N., Warren, G. & Shima, D. T. Golgi clusters and vesicles mediate mitotic inheritance independently of the endoplasmic reticulum. J. Cell Biol. 154, 317–330 (2001).

70. Belevich, I., Joensuu, M., Kumar, D., Vihinen, H. & Jokitalo, E. Microscopy Image Browser: A Platform for Segmentation and Analysis of Multidimensional Datasets. PLOS Biol. 14, e1002340 (2016).

71. Berg, S. et al. ilastik: interactive machine learning for (bio)image analysis. Nat. Methods 16, 1226–1232 (2019).

72. Callister, S. J. et al. Normalization Approaches for Removing Systematic Biases Associated with Mass Spectrometry and Label-Free Proteomics. J. Proteome Res. 5, 277–286 (2006).

73. Huang, T. et al. Combining Precursor and Fragment Information for Improved Detection of Differential Abundance in Data Independent Acquisition*. Mol. Cell. Proteomics 19, 421–430 (2020).

74. Ashburner, M. et al. Gene ontology: tool for the unification of biology. The Gene Ontology Consortium. Nat. Genet. 25, 25–29 (2000).

75. Aleksander, S. A. et al. The Gene Ontology knowledgebase in 2023. Genetics 224, iyad031 (2023).

76. Thomas, P. D. et al. PANTHER: Making genome-scale phylogenetics accessible to all. Protein Sci. 31, 8–22 (2022).

77. Jumper, J. et al. Highly accurate protein structure prediction with AlphaFold. Nature 596, 583– 589 (2021).

78. Varadi, M. et al. AlphaFold Protein Structure Database in 2024: providing structure coverage for over 214 million protein sequences. Nucleic Acids Res. 52, D368–D375 (2024).

79. Jo, S., Kim, T., Iyer, V. G. & Im, W. CHARMM-GUI: A web-based graphical user interface for CHARMM. J. Comput. Chem. 29, 1859–1865 (2008).

80. Huang, J. et al. CHARMM36m: an improved force field for folded and intrinsically disordered proteins. Nat. Methods 14, 71–73 (2017).

81. Jorgensen, W. L., Chandrasekhar, J., Madura, J. D., Impey, R. W. & Klein, M. L. Comparison of simple potential functions for simulating liquid water. J. Chem. Phys. 79, 926–935 (1983).

82. Beglov, D. & Roux, B. Finite representation of an infinite bulk system: Solvent boundary potential for computer simulations. J. Chem. Phys. 100, 9050–9063 (1994).

83. Abraham, M. J. et al. GROMACS: High performance molecular simulations through multi- level parallelism from laptops to supercomputers. SoftwareX 1–2, 19–25 (2015).

84. Hopkins, C. W., Le Grand, S., Walker, R. C. & Roitberg, A. E. Long-Time-Step Molecular Dynamics through Hydrogen Mass Repartitioning. J. Chem. Theory Comput. 11, 1864–1874 (2015).

85. Nosé, S. A molecular dynamics method for simulations in the canonical ensemble. Mol. Phys. 52, 255–268 (1984).

86. Hoover, W. G. Canonical dynamics: Equilibrium phase-space distributions. Phys. Rev. A 31, 1695–1697 (1985).

87. Parrinello, M. & Rahman, A. Polymorphic transitions in single crystals: A new molecular dynamics method. J. Appl. Phys. 52, 7182–7190 (1981).

88. Hess, B. P-LINCS: A Parallel Linear Constraint Solver for Molecular Simulation. J. Chem. Theory Comput. 4, 116–122 (2008).

89. Essmann, U. et al. A smooth particle mesh Ewald method. J. Chem. Phys. 103, 8577–8593 (1995).

90. Humphrey, W., Dalke, A. & Schulten, K. VMD: Visual molecular dynamics. J. Mol. Graph. 14, 33–38 (1996).

91. Gowers, R. J. et al. MDAnalysis : A Python Package for the Rapid Analysis of Molecular Dynamics Simulations MDAnalysis. Proc. 15th Python Sci. Conf. 98–105 (2016) doi:10.25080/Majora-629e541a-00e.

92. Michaud-Agrawal, N., Denning, E. J., Woolf, T. B. & Beckstein, O. MDAnalysis: A toolkit for the analysis of molecular dynamics simulations. J. Comput. Chem. 32, 2319–2327 (2011).

93. Kramer, M. A. Autoassociative neural networks. Comput. Chem. Eng. 16, 313–328 (1992).

94. Paszke, A., et al. PyTorch: An Imperative Style, High-Performance Deep Learning Library. ArXiv abs/1912.0, (2019).

95. Agarap, A. F. M. Deep Learning using Rectified Linear Units (ReLU). ArXiv 2–8 (2019) doi:10.48550/arXiv.1803.08375.

96. Pedregosa, F. et al. Generating the blood exposome database using a comprehensive text mining and database fusion approach. J. Mach. Learn. Res. 12, 2825–2830 (2011).

97. Kumar, S. & Nussinov, R. Close-Range Electrostatic Interactions in Proteins. ChemBioChem 3, 604–617 (2002).

