## Supplemental Figures for "Control of Golgi- V-ATPase through Sac1-dependent co-regulation of PI(4)P and cholesterol"

**Supplementary Figure 1.**

**a-c)** Graph showing the total PI(3)P (**a**) PI(4,5)P2 (**b**) and PI(3,5)P2 (**c**) levels (pmol/x$10^6$ cells) ±S.E.M. of Sac1-degron A431 cells treated with DMSO for 2 h or IAA for 0.5, 1 or 2 h as measured by PRMC-MS. n = 3 biological replicates. One-way ANOVA with Dunnett's multiple comparisons test. n.s. = non-significant, * p<0.05, ** p < 0.01, *** p<0.005.

**d)** Bar graphs depicting the average PI(4)P, PI(3)P, PI(4,5)P2 and PI(3,5)P2 levels (pmol/x$10^6$ cells) ±S.E.M. per molecular species of Sac1-degron A431 cells treated with DMSO for 2 h or IAA for 0,5, 1 or 2 h as measured by PRMC-MS.

**e-h)** Bar graph depicting the average levels of PI(4)P (**e**), PI(3)P (**f**), PI(4,5)$P_2$ (**g**), PI(3,5)$P_2$ (**h**) (pmol/x$10^6$ cells) ±S.E.M. of Sac1-degron A431 cells treated with DMSO or IAA for 4 h as measured by PRMC-MS. n = 3 biological replicates. Student's t-test. ** p < 0.01, *** p < 0.005, **** p < 0.001.

**i, j)** Representative confocal images of Sac1-degron HEK293A cells (**i**) and A549 cells (**j**) treated for, respectively, 3 or 4 h with DMSO or IAA and stained for TGN46.

**Supplementary Figure 2.**

**a)** Volcano plot showing the up and downregulated proteins by comparing the proteome profile of 3 h IAA treated Sac1-degron A431 cells to DMSO control.

**b)** Relative mRNA levels of TGN46 and B4GALT1 of Sac1-degron A431 cells treated with IAA for 4 h with DMSO or IAA. Data from 5 biological replicates. Two-way ANOVA with *Šídák* multiple comparisons test. n.s. = non-significant.

**c)** Western blot analysis of Sac1-degron A431 cells treated with IAA for 0 to 4 h with or without bafilomycin A (BafA1). Blotted for miniIAA7 (Sac1), TGN46 and Golgin-97.

**d)** Western blot analysis of Sac1-degron A431 cells treated for 0 to 8 h with IAA and blotted for TGN46. Black arrow indicates mature TGN46, and red arrow indicates the immature form of TGN46.

**e)** Western blot analysis of Sac1-degron A431 cells treated for 8 h with DMSO or IAA and with or without PNGaseF digestion. Blotted for TGN46.

**f)** Western blot analysis of Sac1-degron A431 cells treated for 8 h with DMSO or IAA and with or without α2-3,6,8,9 Neuraminidase A digestion. Blotted for TGN46.

**Supplementary Figure 3.**

**a)** Representative confocal images of Sac1-degron A431 cells transiently transfected with GalT-RpHLuorin2 (green) and co-stained with Golgin-97 (magenta).

**b)** Representative pH calibration curve of Sac1-degron A431 cells transiently transfected with GalT-RpHLuorin2. Calibration buffers of pH 4.5, 5.5, 6.5 and 7.5 were used. Y-axis shows lifetime (ns), and x-axis shows the pH.

**c)** Representative phasor plots of FLIM imaging of Sac1-degron A431 cells transiently transfected with GALT1-RpHLuorin2 treated with DMSO or IAA for 1 and 3 h. Dotted red line shows the shift in lifetime.

**d)** Western blot analysis of Sac1-degron A431 cell lysates treated for 0, 1, 2 or 4 h with IAA. Stained for ATP6$V_0$d1 or ATP6$V_1$A.

**e)** Western blot analysis of membrane and cytosolic fractions of A431 Sac1-degron cells treated for 2 h with DMSO or IAA. Blotted for ATP6$V_1$A and ATP6$V_0$d1.

**f)** Graph depicting the percentage of total ATP6$V_1$A subunit in the cytosol. Data from 4 independent experiments. Paired Student's t-test n.s. = non-significant.

**g)** Representative confocal images of Sac1-degron A549 cells treated with DMSO or IAA for 2 h and stained for ATP6$V_0$a2 (red), GM130 (green) and LAMP2 (cyan).

**Supplementary Figure 4.**

**a)** Representative images of Sac1-degron A431 cells transfected with mCherry-P4M-SidM and treated with IAA for 0, 30, 60 or 120 min. mCherry-P4M-SidM intensity is converted into a heat map (0–6600 a.u.).

**b)** Representative widefield images of Sac1-degron A431 cells treated with DMSO for 4 h (0) or IAA for 1, 2 or 4 h and stained with PI(4)P antibody.

**c)** Violin plot depicting the average PI(4)P staining intensity from (**b**). n = 80 images. Representative data from one of three independent experiments. One-way ANOVA with Dunnett's multiple comparisons test. **** $p < 0.001$.

**d)** Representative widefield images of Sac1-degron A431 cells treated with DMSO 4 h (0) or IAA for 1, 2 or 4 h and stained with mNeonGreen-ALOD4.

**e)** Violin plot depicting the average intensity of mNeonGreen-ALOD4 calculated from (**d**). n = 50 images. Representative data from one of three independent experiments. One-way ANOVA with Dunnett's multiple comparisons test. *** $p<0.005$, **** $p<0.001$.

**f)** Representative images of Sac1-degron A431 cells stained for free cholesterol with Filipin and treated with IAA for 0 to 4 h.

**g)** Violin plot depicting the distribution of the average Filipin intensity calculated from (**f**). n > 40 images. Representative data from one of three independent experiments. One-way ANOVA with Dunnett's multiple comparisons test. **** $p<0.001$.

**h)** Bar graph depicting the average content (µg/mg protein) of free cholesterol (FC)±S.E.M. in Sac1-degron A431 cells treated with IAA for 0 to 4 h as measured by HPTLC. Data from 3 independent experiments. One-way ANOVA with Dunnett's multiple comparisons test. n.s. = non-significant.
**i)** Representative confocal images of Sac1-degron A431 treated for 2 h with DMSO or IAA and stained for filipin (magenta) and TMEM192 (yellow).
**j)** Bar graph showing normalized Filipin intensity in the lysosomes from (**i**). n = 21 – 31 images, data from 3 independent experiments. Mann-Whitney U test. * p<0.05.
**k)** Bar graph showing the average alkyne-oleic acid incorporation into cholesteryl esters ±S.E.M. Alkyne-OA was loaded during the last 30 min of 2 h DMSO or IAA treatment in Sac1-degron A431 cells. n = 4 biological replicates, Student's t-test. **** p < 0.001.
**l)** Bar graph depicting the average content (µg/mg protein) of cholesterol ester (CE) ±S.E.M. in Sac1-degron A431 cells treated with IAA for 0 to 4 h as measured by HPTLC. Data from 3 independent experiments. One-way ANOVA with Dunnett's multiple comparisons test. n.s. = non-significant, * p < 0.05, ** p < 0.01.

**Supplementary Figure 5.**
**a)** Western blot analysis of sucrose gradient fractionations (sucrose concentration from 0.3M (fraction 1) to 2.1M (loading fractions 9-12)) of Sac1-degron A431 cells treated with or without methyl-β cyclodextrin (MβCD) and blotted for ATP6$V_0$a2.
**b)** Representative confocal images of Sac1-degron A549 cells treated with DMSO, IAA, IAA with cholesterol or IAA with PI4K inhibitors for 2 h. Stained for GM130 (green), B4GALT1 (magenta) and TGN46 (red).
**c)** Bar graph depicting the percentage (mean ±S.E.M) of cells with intact, partially fragmented, or fully fragmented Golgi in (**b**). n = 101 - 139 cells.
**d)** Violin plot showing the average TGN46 intensity in the Golgi area (GM130) of (**b**). n = 102 – 118 cells. Kruskall-Wallis with Dunnet's multiple correction test. n.s. = non-significant, **** p<0.001.

**Supplementary Figure 6.**
**a)** The four clusters identified by BGMM have similar populations indicative of balanced sampling.
**b)** Root Mean Square Deviation (RMSD) of heavy atoms of all structures in the same cluster, with the cluster mean being ~5 Å, indicative of high structural similarity within each cluster. The error bars are the standard error computer over 32 independent datasets composed of 8 independent replicas with 4 different lipid compositions.

**c)** Representative side view of the $V_0$ region from each of the four clusters. In each cluster, it is chosen as the structure with the smallest distance from the mean of the Gaussian in the BGMM.

**d)** Occupational density maps of the lipids in the vicinity of the $V_0$ region (occupancy level 45%). Left: a representative active conformation (10% PI(4)P (orange), 20% cholesterol (green)). Right: a representative inactive conformation (20% PI(4)P).

**Supplementary Figure 7.**

**a)** Representative fluorescence micrographs of wild type hESCs differentiated into trophoblasts and stained for the trophoblast markers hCGβ, GATA3 and Dab2.

**b)** Western blot analysis of hESC and TB lysates blotted for hCGβ.

**c)** Western blot analysis of Sac1-degron STB treated for 0, 2, 4, 8 or 15 h with IAA. Blotted for miniIAA7 (Sac1), TGN46 and B4GALT1.

**d)** Bar graph depicting the TGN46 or B4GALT1 degradation efficiency calculated from (**c**). Data from 3 biological replicates. One-way ANOVA with Dunnett's multiple comparisons test. n.s. = non-significant, * $p<0.05$, ** $p<0.01$, *** $p<0.005$, **** $p<0.0001$.

**e)** Representative confocal images of Sac1-degron TBs treated with DMSO or IAA for 1 to 2 h and stained for PI(4)P (green) and TGN46 (magenta).

**f)** Representative confocal images of Sac1-degron TBs treated for 4 h with DMSO or IAA and stained for Golgin-97 (green) TMEM192 (yellow) and filipin (magenta).

**g)** Violin plot showing normalized filipin intensity in the lysosomes of (**f**) n = 24 – 26 cells, data from 3 independent experiments. Mann-Whitney U test. n.s. = non-significant

**Extended Data 1.**

Quantification of Golgi morphological changes from TEM images of Sac1-degron A431 cells treated with DMSO or IAA for 2 h.

**Extended Data 2.**

Results of the proteomics analysis of Sac1-degron A431 cells treated with DMSO or IAA for 3 h.

**Movie 1, 2**

Partial Least Squares based Discriminant Analysis (PLS-DA) of the four clusters. The movies highlight the largest structural changes of the $V_0$ region as it moves from clusters 1, 3, and 4 (impairing assembly) to cluster 2 (favoring assembly) from top (**Movie 1**) and side view (**Movie 2**). Along the direction shown in the movies, the a2 head (green) moves away from the d1 subunit (magenta) and toward the membrane, preventing potential structural clashes of the a2 arm (yellow) and the a2 head with the $V_1$ region.

### Suppl. Fig. 1

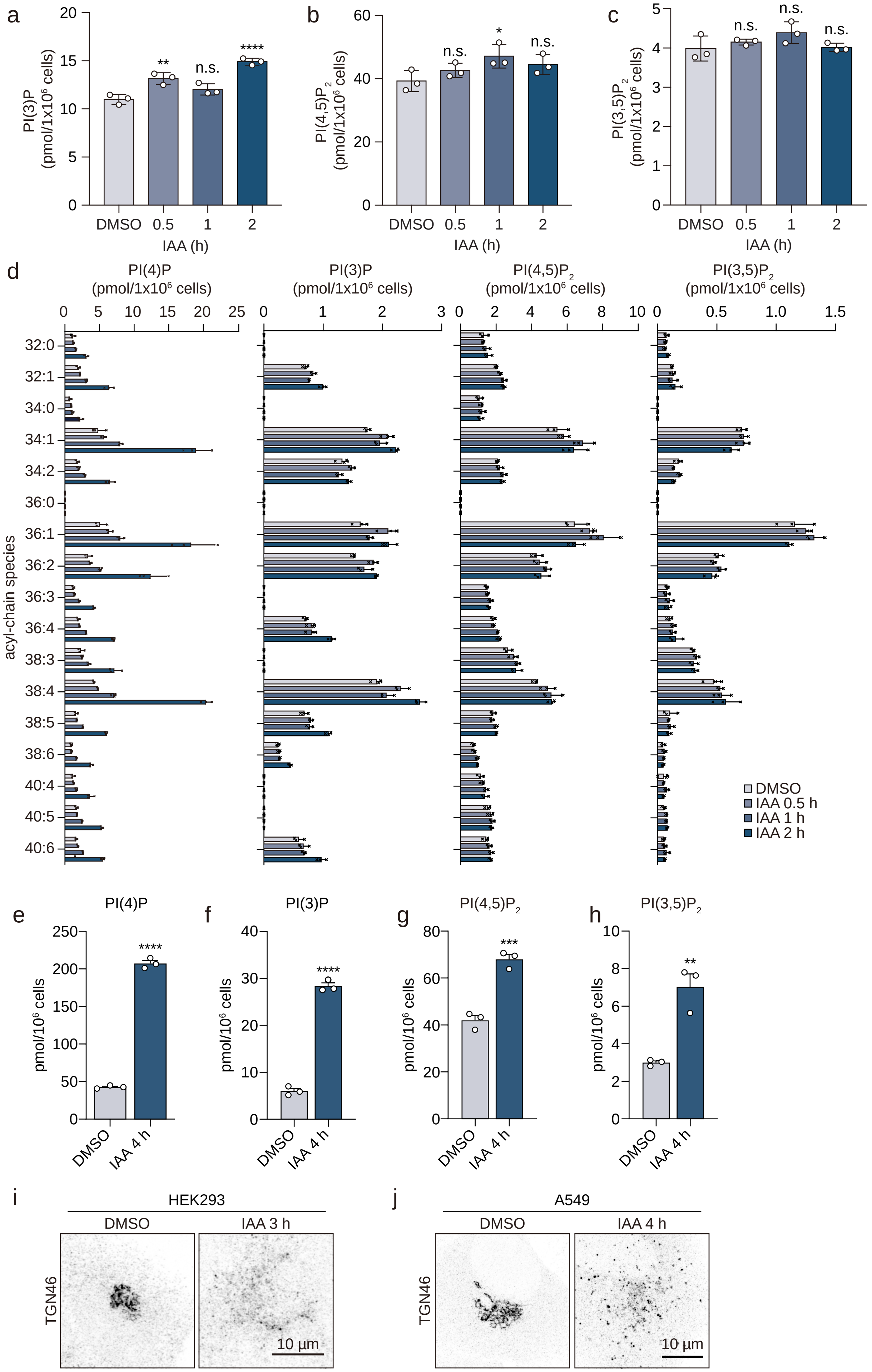

a
PI(3)P (pmol/1x10^6 cells)
20
15
10
5
0
**
n.s.
****
DMSO
0.5
1
2
IAA (h)
b
PI(4,5)P2 (pmol/1x10^6 cells)
60
40
20
0
n.s.
*
n.s.
DMSO
0.5
1
2
IAA (h)
c
PI(3,5)P2 (pmol/1x10^6 cells)
5
4
3
2
1
0
n.s.
n.s.
n.s.
DMSO
0.5
1
2
IAA (h)
d
PI(4)P (pmol/1x10^6 cells)
0 5 10 15 20 25
PI(3)P (pmol/1x10^6 cells)
0 1 2 3
PI(4,5)P2 (pmol/1x10^6 cells)
0 2 4 6 8 10
PI(3,5)P2 (pmol/1x10^6 cells)
0 0.5 1.0 1.5
acyl-chain species
32:0
32:1
34:0
34:1
34:2
36:0
36:1
36:2
36:3
36:4
38:3
38:4
38:5
38:6
40:4
40:5
40:6
DMSO
IAA 0.5 h
IAA 1 h
IAA 2 h
e
PI(4)P
pmol/10^6 cells
250
200
150
100
50
0
****
DMSO
IAA 4 h
f
PI(3)P
pmol/10^6 cells
40
30
20
10
0
****
DMSO
IAA 4 h
g
PI(4,5)P2
pmol/10^6 cells
80
60
40
20
0
***
DMSO
IAA 4 h
h
PI(3,5)P2
pmol/10^6 cells
10
8
6
4
2
0
**
DMSO
IAA 4 h
i
HEK293
DMSO
IAA 3 h
TGN46
10 µm
j
A549
DMSO
IAA 4 h
TGN46
10 µm

### Suppl. Fig. 2

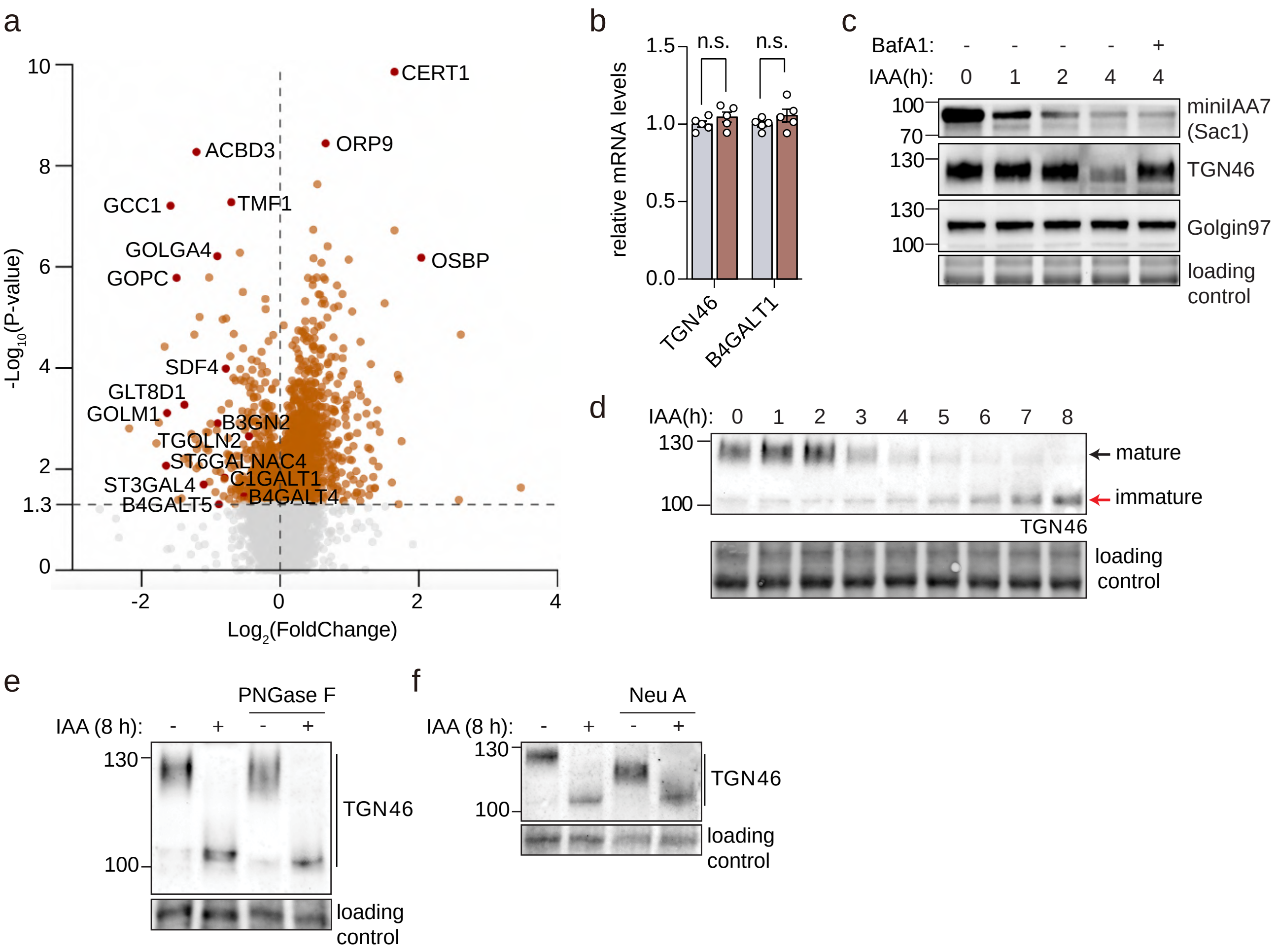
a
CERT1
ACBD3
ORP9
GCC1
TMF1
GOLGA4
OSBP
GOPC
SDF4
GLT8D1
GOLM1
B3GN2
TGOLN2
ST6GALNAC4
C1GALT1
ST3GAL4
B4GALT4
B4GALT5
-Log10(P-value)
Log2(FoldChange)
10
8
6
4
2
1.3
0
-2
0
2
4
b
relative mRNA levels
n.s.
n.s.
1.5
1.0
0.5
0.0
TGN46
B4GALT1
c
BafA1: - - - - +
IAA(h): 0 1 2 4 4
100
70
130
130
100
miniIAA7
(Sac1)
TGN46
Golgin97
loading
control
d
IAA(h): 0 1 2 3 4 5 6 7 8
130
100
mature
immature
TGN46
loading
control
e
PNGase F
IAA (8 h): - + - +
130
100
TGN46
loading
control
f
Neu A
IAA (8 h): - + - +
130
100
TGN46
loading
control

### Suppl. Fig. 3

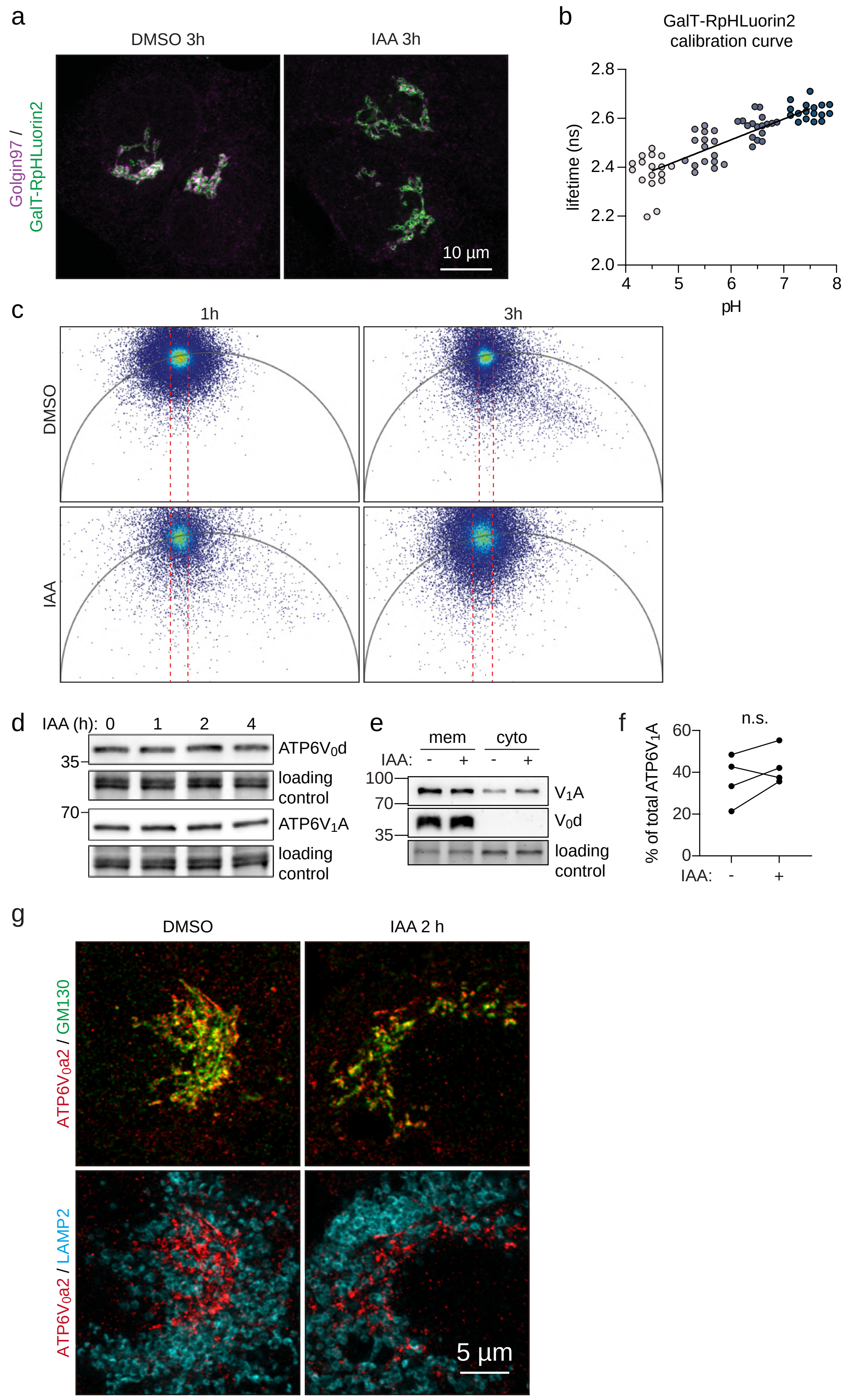

a
DMSO 3h
IAA 3h
Golgin97 / GalT-RpHLuorin2
10 µm
b
GalT-RpHLuorin2 calibration curve
lifetime (ns)
2.8
2.6
2.4
2.2
2.0
4
5
6
7
8
pH
c
1h
3h
DMSO
IAA
d
IAA (h): 0 1 2 4
35
ATP6V0d
loading control
70
ATP6V1A
loading control
e
mem
cyto
IAA: - + - +
100
70
35
V1A
V0d
loading control
f
n.s.
% of total ATP6V1A
60
40
20
0
IAA: - +
g
DMSO
IAA 2 h
ATP6V0a2 / GM130
ATP6V0a2 / LAMP2
5 µm

### Suppl. Fig. 4

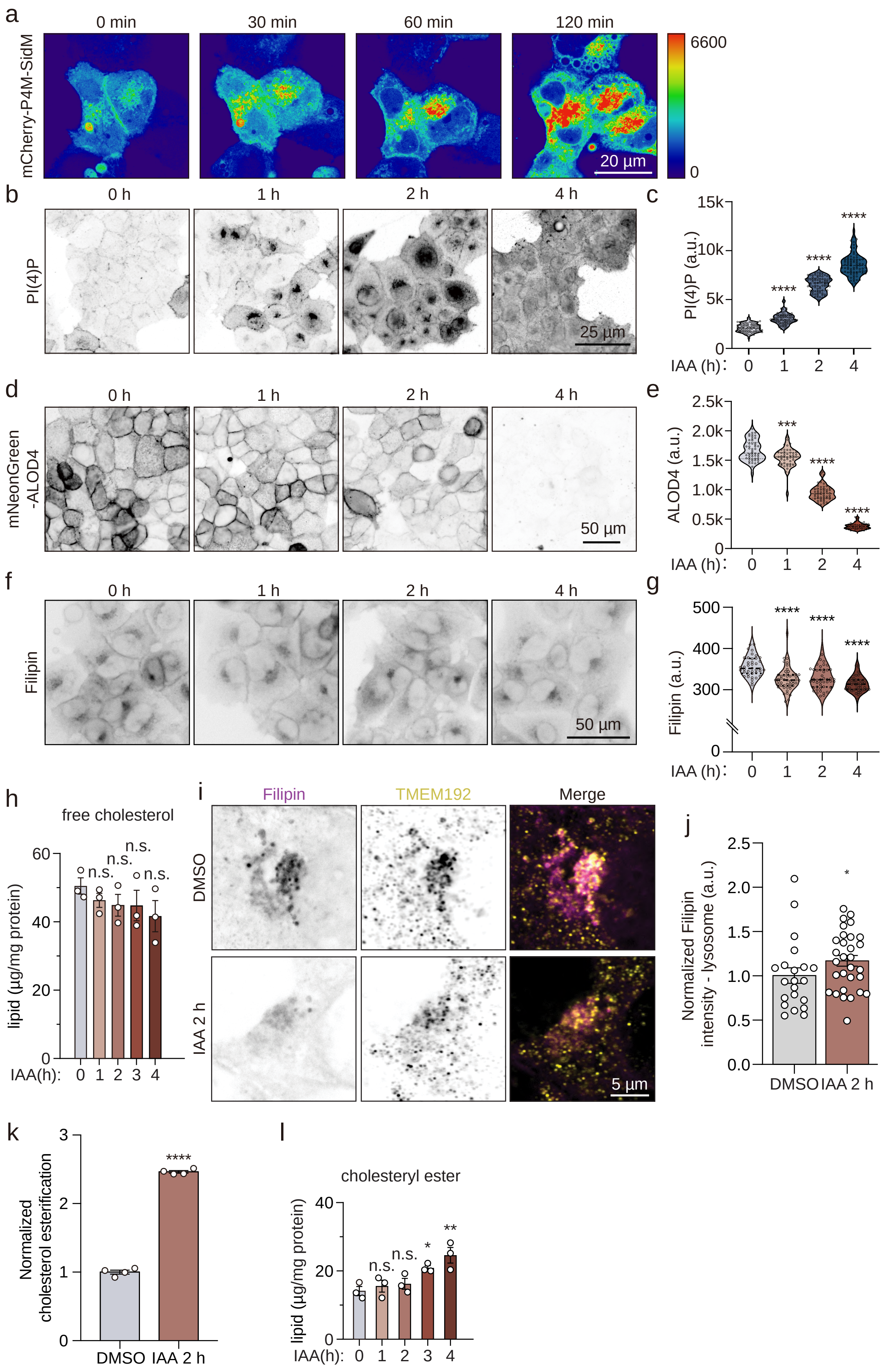

a
0 min
30 min
60 min
120 min
mCherry-P4M-SidM
6600
0
20 µm
b
0 h
1 h
2 h
4 h
PI(4)P
25 µm
c
PI(4)P (a.u.)
IAA (h):
d
mNeonGreen -ALOD4
50 µm
e
ALOD4 (a.u.)
f
Filipin
50 µm
g
Filipin (a.u.)
h
free cholesterol
lipid (µg/mg protein)
n.s.
IAA(h):
i
Filipin
TMEM192
Merge
DMSO
IAA 2 h
5 µm
j
Normalized Filipin intensity - lysosome (a.u.)
DMSO
IAA 2 h
k
Normalized cholesterol esterification
l
cholesteryl ester

### Suppl. Fig. 5

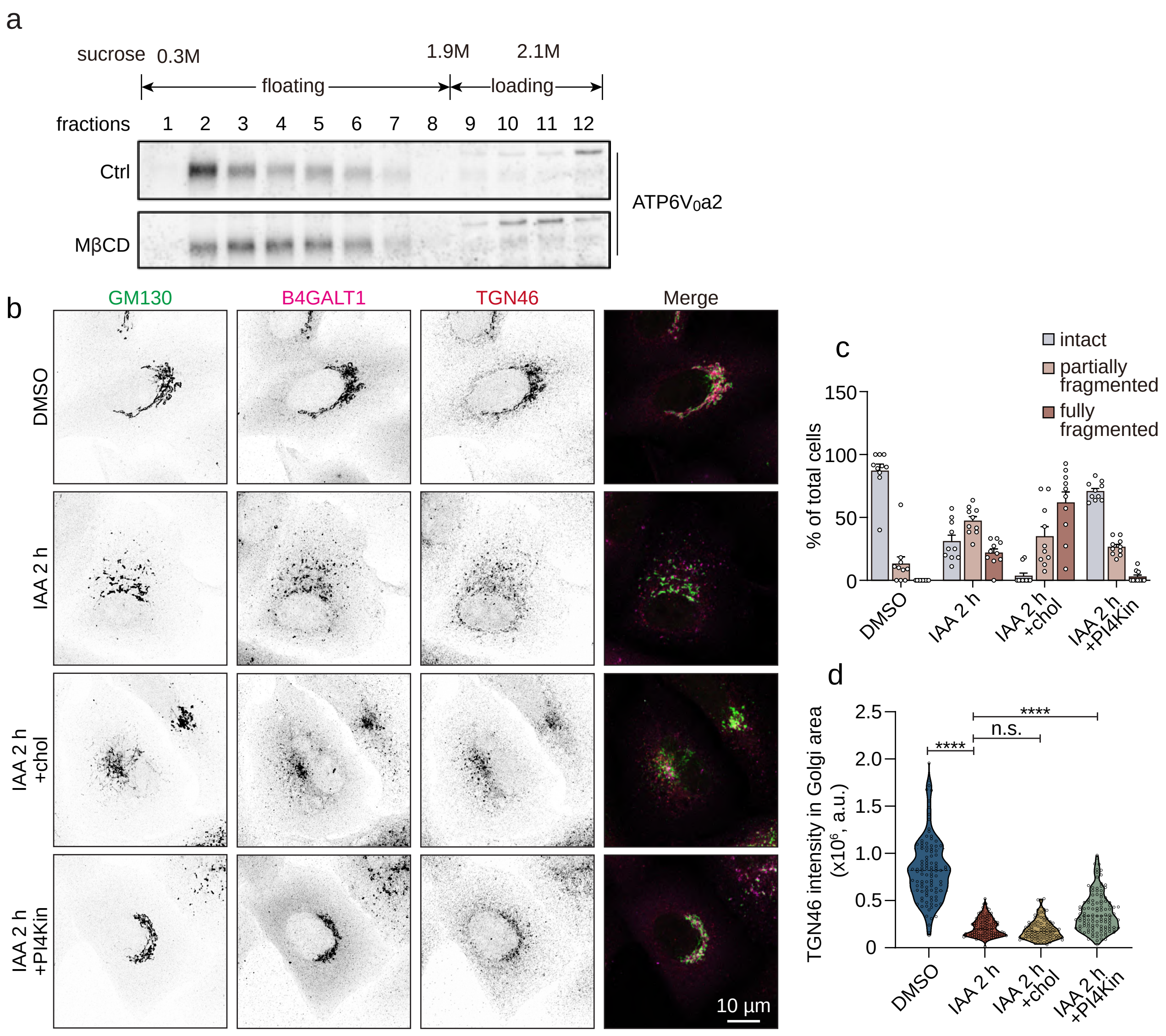
a
sucrose 0.3M
1.9M
2.1M
floating
loading
fractions 1 2 3 4 5 6 7 8 9 10 11 12
Ctrl
MβCD
ATP6V0a2
b
GM130
B4GALT1
TGN46
Merge
DMSO
IAA 2 h
IAA 2 h +chol
IAA 2 h +PI4Kin
10 µm
c
intact
partially fragmented
fully fragmented
% of total cells
150
100
50
0
DMSO
IAA 2 h
IAA 2 h +chol
IAA 2 h +PI4Kin
d
TGN46 intensity in Golgi area (x10^6, a.u.)
2.5
2.0
1.5
1.0
0.5
0
****
n.s.
****
DMSO
IAA 2 h
IAA 2 h +chol
IAA 2 h +PI4Kin

### Suppl. Fig. 6

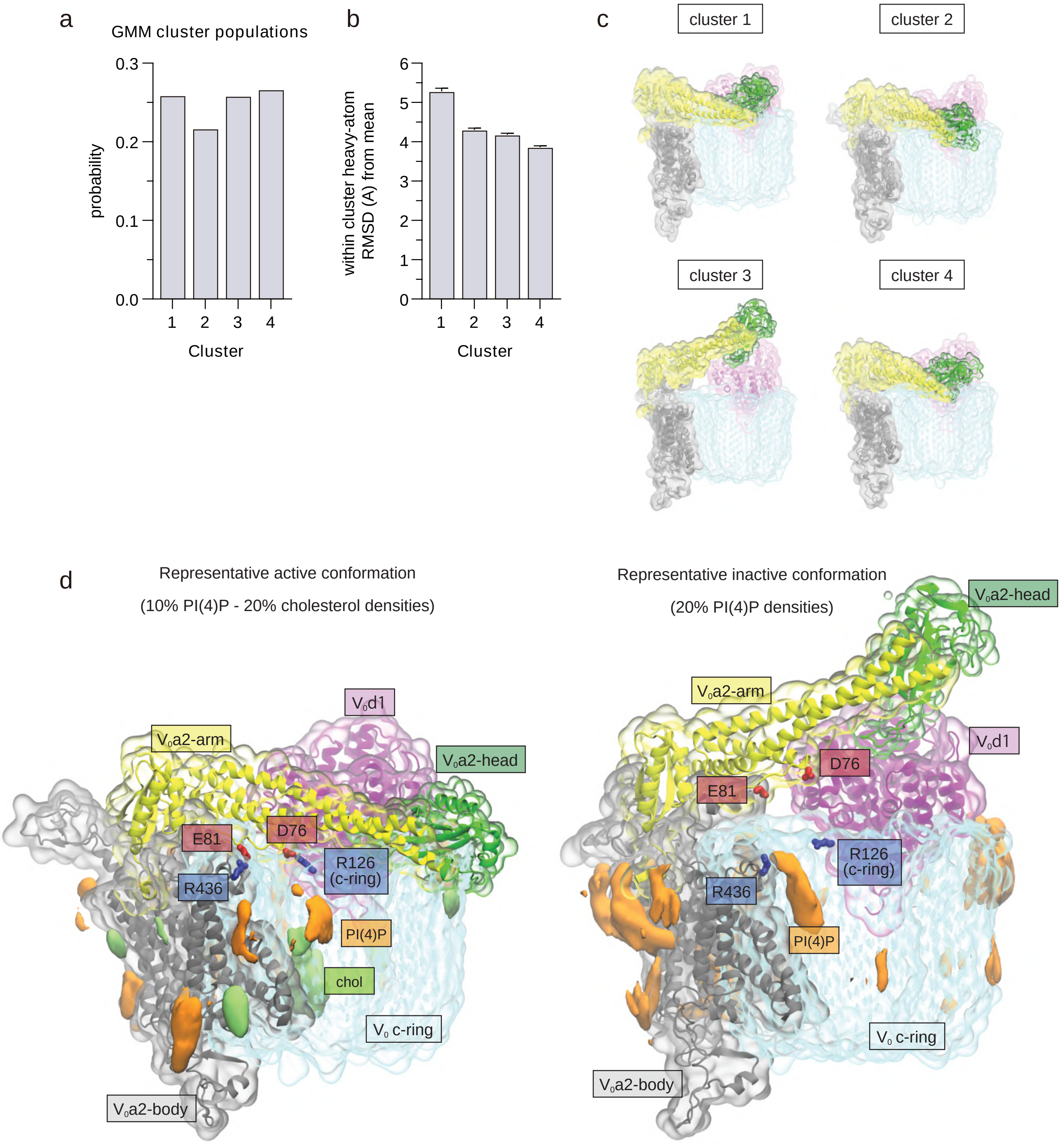

a
GMM cluster populations
probability
0.3
0.2
0.1
0.0
1
2
3
4
Cluster
b
within cluster heavy-atom RMSD (A) from mean
6
5
4
3
2
1
0
1
2
3
4
Cluster
c
cluster 1
cluster 2
cluster 3
cluster 4
d
Representative active conformation
(10% PI(4)P - 20% cholesterol densities)
Representative inactive conformation
(20% PI(4)P densities)
V₀d1
V₀a2-arm
V₀a2-head
E81
D76
R126 (c-ring)
R436
PI(4)P
chol
V₀ c-ring
V₀a2-body
V₀a2-head
V₀a2-arm
V₀d1
D76
E81
R126 (c-ring)
R436
PI(4)P
V₀ c-ring
V₀a2-body

### Suppl. Fig. 7

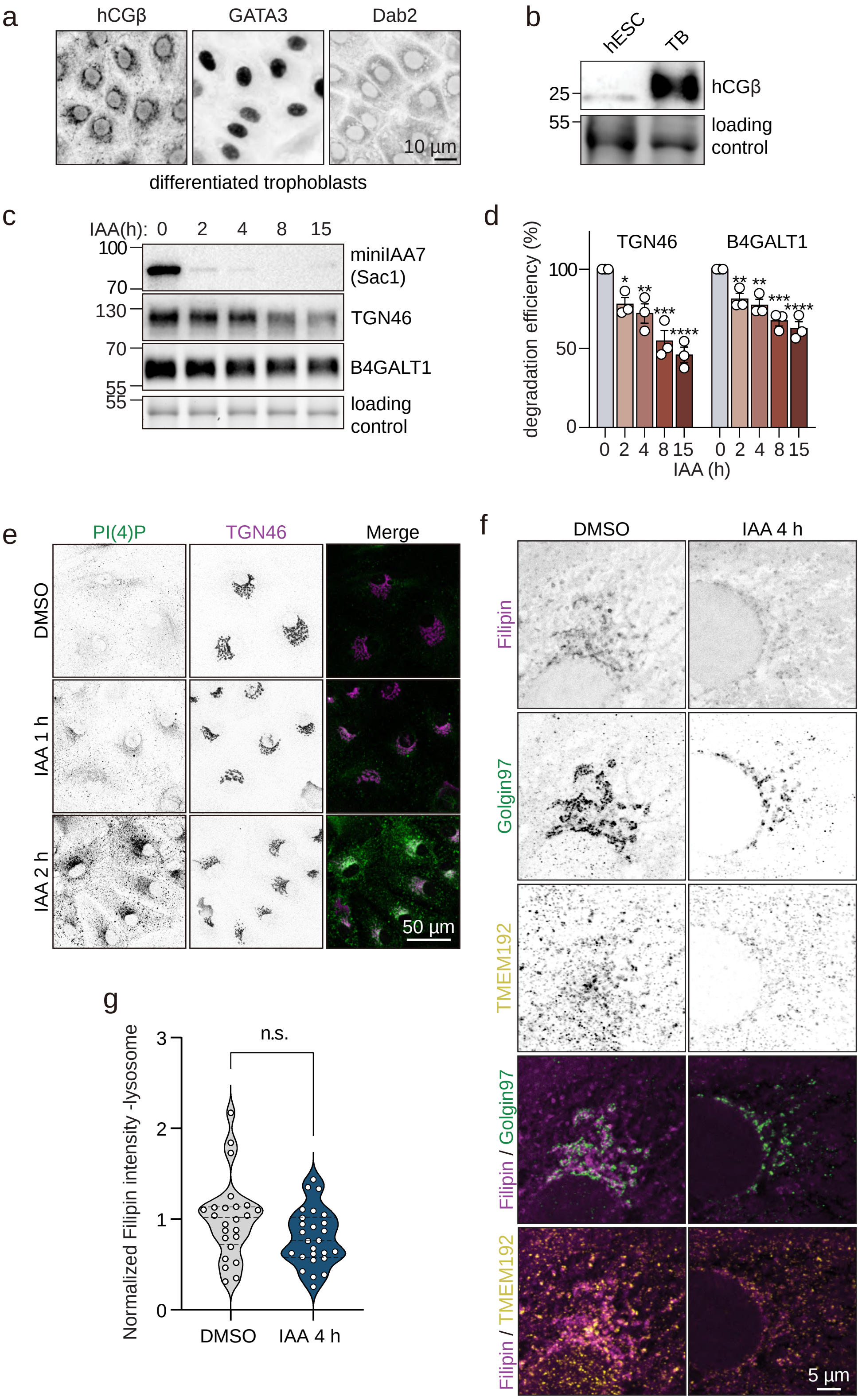
a
hCGβ
GATA3
Dab2
10 µm
differentiated trophoblasts
b
hESC
TB
25
55
hCGβ
loading control
c
IAA(h): 0 2 4 8 15
100
70
130
70
55
55
miniIAA7 (Sac1)
TGN46
B4GALT1
loading control
d
degradation efficiency (%)
TGN46
B4GALT1
100
50
0
0 2 4 8 15 0 2 4 8 15
IAA (h)
e
PI(4)P
TGN46
Merge
DMSO
IAA 1 h
IAA 2 h
50 µm
f
DMSO
IAA 4 h
Filipin
Golgin97
TMEM192
Filipin / Golgin97
Filipin / TMEM192
5 µm
g
Normalized Filipin intensity -lysosome
n.s.
3
2
1
0
DMSO
IAA 4 h
